# An insular cortex to lateral amygdala pathway in fear learning

**DOI:** 10.1101/2023.01.17.524434

**Authors:** Shriya Palchaudhuri, Denys Osypenko, Olexiy Kochubey, Ralf Schneggenburger

## Abstract

During fear learning, associations between an aversive stimulus (the US), and a sensory cue (CS) are formed at specific brain synapses. Nevertheless, how US information is transmitted to brain areas involved in value processing, like the amygdala, is still elusive. Using optogenetics, *in-vivo* Ca^2+^ imaging, and circuit tracing, we investigate the role of the posterior insular cortex (pInsCx) and relevant output pathways of this cortical area in fear learning. Optogenetic suppression of US-signaling in pInsCx principal neurons compromises auditory-cued fear learning. The pInsCx makes a robust glutamatergic synapse in the lateral amygdala (LA), which undergoes long-term potentiation after fear learning, and transmits US-information to a sub-population of LA neurons. Suppressing US-signaling in LA-projectors recapitulates the fear learning deficits observed after silencing pInsCx principal neurons. Thus, the pInsCx, via a plastic output synapse, transmits US-information to the LA and critically contributes to the formation of auditory-cued fear memories.

## Introduction

Animals and humans must evaluate signs of dangers in their environment and react with appropriate behavior, to ensure survival. For this reason, robust mechanisms of aversively-motivated associative learning, also called fear learning, have evolved (LeDoux, 2000; Janak and Tye, 2015; Feinberg and Mallatt, 2017; Seymour, 2019). During fear learning, animals learn to associate an innocuous sensory cue, like a tone (the conditioned stimulus, CS), with a harmful event like a nociceptive stimulus (unconditioned stimulus, US). The amygdala, and especially the lateral amygdala (LA) has an important role in associative learning, and there is broad agreement that integration of US- and CS - information, and long-term plasticity at synapses that code for CS information, is necessary for fear learning (LeDoux, 2000; Maren, 2001; Sigurdsson et al., 2007; Herry and Johansen, 2014; Tovote et al., 2015). The LA receives input from non-lemniscal auditory thalamic nuclei (LeDoux et al., 1990; Barsy et al., 2020), as well as from ventral, higher-order areas of the auditory cortex (Romanski and LeDoux, 1993; Shi and Cassell, 1997; McDonald, 1998; Dalmay et al., 2019). Fear learning has been shown to induce signs of long-term potentiation (LTP) at the input synapses from the thalamus (Rogan and LeDoux, 1995; McKernan and Shinnick-Gallagher, 1997; Rumpel et al., 2005; Clem and Huganir, 2010; Namburi et al., 2015) and at synapses from the auditory cortex to the LA (Kim and Cho, 2017). Furthermore, optogenetic induction of LTP, or LTD at both input synapses *in-vivo* strongly modulates the fear state of rats (Nabavi et al., 2014), and blocking a postsynaptic form of LTP in the LA compromises auditory-cued fear learning (Rumpel et al., 2005). Thus, there is good evidence that long-term plasticity at either the thalamic input alone, or at both the thalamic and auditory cortex inputs to the LA together, is critically involved in basic forms of auditory-cued fear learning.

Despite these findings on the synaptic *substrate* for LTP, it has remained elusive which synapses drive action potential (AP) firing of LA neurons during the footshock, to cause associative plasticity in the LA that underlies fear learning (Herry and Johansen, 2014; Johansen et al., 2014). *In-vivo* Ca^2+^ imaging and cFos labeling have shown that sub-populations of LA neurons, or in some studies BA neurons, are activated by footshocks (Gore et al., 2015; Grewe et al., 2017; Zhang and Li, 2018). More specifically, it has been shown that a depolarizing drive of LA principal neurons, in addition to neuromodulatory influences, causes associative synaptic plasticity in LA neurons, which in turn contributes to auditory-cued fear learning (Johansen et al., 2010; Herry and Johansen, 2014; Johansen et al., 2014). Nevertheless, the presynaptic source of this depolarizing drive for associative plasticity in the LA has not been identified. Recent studies have shown that the inputs from the auditory cortical - and thalamic areas can themselves carry US - information (Dalmay et al., 2019; Barsy et al., 2020; Taylor et al., 2021). However, the implications of these findings for models of associative learning are not well understood. It is likely that further synaptic afferents carry footshock information to the LA, and to the basal amygdala (BA).

The insular cortex is an interesting candidate brain area when considering pathways that might carry US information to the basolateral amygdala (BLA) complex. In rodents, the insular cortex lies ventral to the motor - and secondary somatosensory cortices (S2), and can be separated into an anterior, middle, and posterior part (Gehrlach et al., 2020). The posterior insular cortex (pInsCx) is known to code for nociceptive stimuli in humans (Apkarian et al., 2005; Coghill, 2020) and in rats (Rodgers et al., 2008), and neuroanatomical studies in rats and monkeys have revealed a projection from the pInsCx to the LA (Mufson et al., 1981; McDonald, 1998; Shi and Cassell, 1998). Moreover, recent studies showed that the pInsCx codes for prolonged aversive states in mice (Gehrlach et al., 2019), and a cortical area that closely overlaps with the pInsCx codes for skin temperature (Bokiniec et al., 2022; Vestergaard et al., 2022), a sensory modality that has close links with nociception (Basbaum and Jessell, 2013). Nevertheless, there is still no agreement on the role of the pInsCx in fear learning. An earlier lesion study in rats has found evidence for a role of the pInsCx in fear learning (Shi and Davis, 1999), while others have not (Brunzell and Kim, 2001; Lanuza et al., 2004). More recent studies using pharmacological inactivation have described roles of the pInsCx in safety learning (Foilb et al., 2016), and in fear memory consolidation (Casanova et al., 2016; de Paiva et al., 2021). Also, the notion of a pInsCx to LA connection has not been confirmed in recent anatomical studies in mice, which found outputs from the pInsCx to the central amygdala (CeA), but only weak ones to the LA (Gehrlach et al., 2019; Gehrlach et al., 2020). Thus, the question whether the pInsCx processes US-information during fear learning, and sends it to the LA or to other components of the BLA complex, has remained unanswered.

Here, we have used optogenetic - and circuit-mapping techniques, to investigate the role of the pInsCx in fear learning, and the function of a pInsCx to LA synapse in this process. We show that footshock-induced activity of pInsCx principal neurons is necessary for the efficient formation of an auditory-cued fear memory. Given this role of the pInsCx in fear learning, we mapped its output projections and found that the pInsCx provides a robust glutamatergic connection to the anterior LA, which undergoes LTP following fear learning. Refining the optogenetic silencing approach to LA - projectors in the pInsCx recapitulates the deficits in fear memory formation observed after more widespread silencing of pInsCx principal neurons. Finally, *in-vivo* Ca^2+^ imaging combined with axon silencing revealed that the pInsCx - LA excitatory synapse drives US-responses in a sub-population of LA neurons. Thus, our work identifies the pInsCx as a cortical area that is upstream of the LA, involved in transmitting behaviorally-relevant US-information to the BLA complex during fear learning.

## Results

### Footshock-driven activity of pInsCx principal neurons is necessary for fear learning

The posterior insular cortex (pInsCx) is a candidate brain area for transmitting nociceptive information to the amygdala (McDonald, 1998; Shi and Cassell, 1998; Rodgers et al., 2008; Gehrlach et al., 2019, and see Introduction). Therefore, we investigated whether footshock-driven activity in the pInsCx is a necessary component for auditory-cued fear learning, by performing optogenetic silencing experiments. For this, we employed the light-gated H+ pump Archaerhodopsin (Arch; bilateral injection of AAV1:CAG:FLEX:Arch-eGFP), and CaMKII^Cre^ mice to limit the expression of Arch to principal neurons. In the same surgery, optic fibers were placed above the injection sites (Figure 1A, B). Mice in a control group received bilateral injections of an AAV driving the expression of eGFP, but underwent otherwise identical procedures as the Arch-expressing mice (see Materials and Methods). Control experiments in slices showed that activation of Arch strongly hyperpolarizes principal neurons in the pInsCx, and blocks AP firing evoked by current injections (Figure S1A-G). Furthermore, *in-vivo* optrode recordings in the pInsCx showed that activation of Arch by green light significantly reduced footshock-driven neuronal activity (Figure S1H-K; N=2 mice, p=0.034, W=54; Wilcoxon’s test). Post-hoc analyses of injection sites and optic fiber placements showed that Arch - mediated inhibition was directed to the pInsCx - GI and DI; it remains possible that principal neurons in the ventral part of the S2, close to the S2 - pInsCx border, were also silenced (Figure 1B; Figure S1L, M).

**Figure 1.**
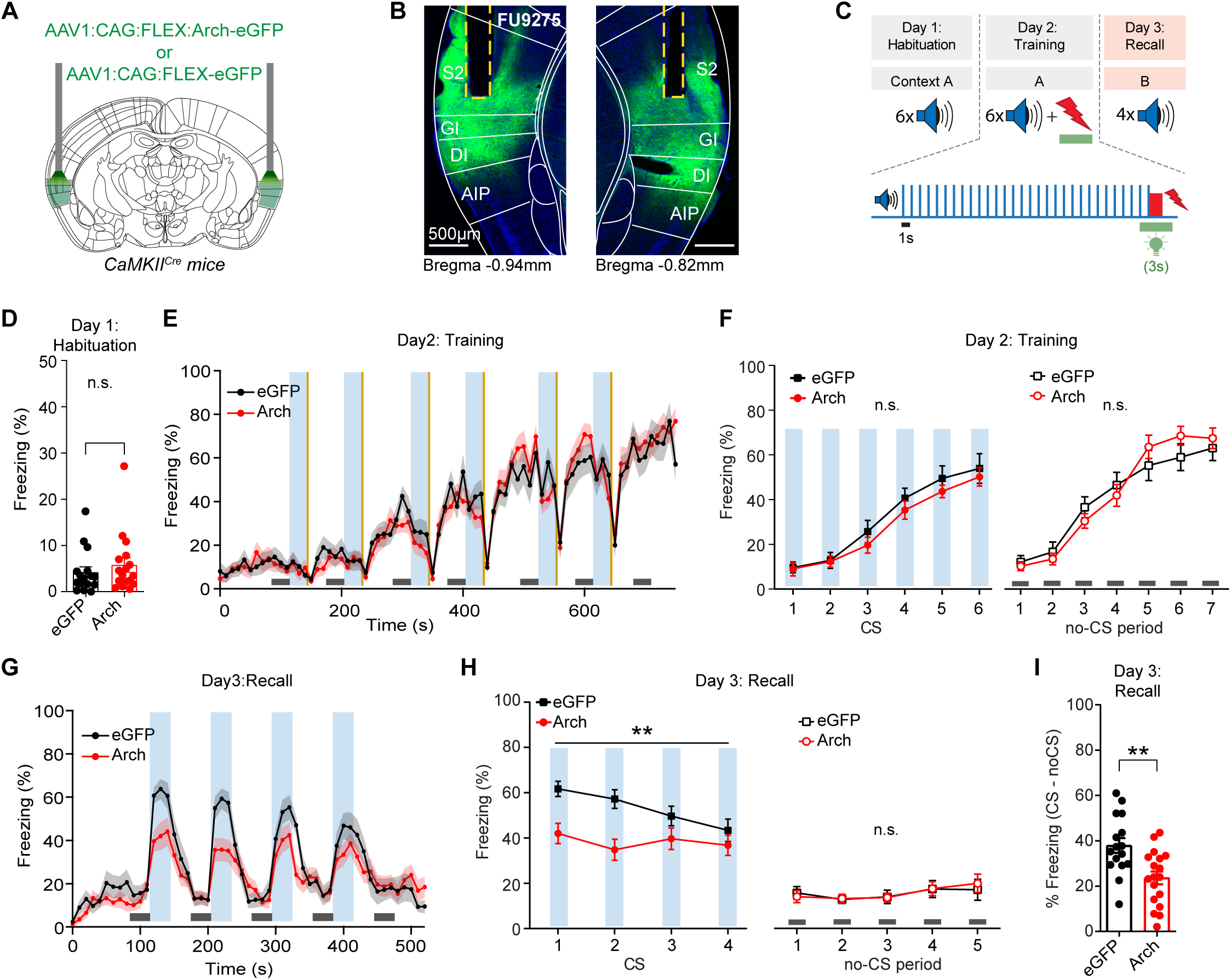
Optogenetic suppression of footshock-driven activity of pInsCx principal neurons impairs fear memory formation. (A) Scheme of the experimental approach. (B) A representative fluorescence image showing the expression of Arch-eGFP in the pInsCx, and the placement of optic fibres. Brain atlas overlays were from (Franklin and Paxinos, 2016; see also Table 1 for abbreviations of brain structures). (C) Scheme of the behavioral protocol. During the training day, brief (3s) pulses of green light (561 nm) centered around every footshock presentation were applied. (D) Freezing during the CS blocks on the habituation day for Arch - and eGFP - expressing mice (N = 17 and 16; red and black data points; p = 0.292, Mann Whitney U test). (E) Time-resolved percent of time spent freezing for the training session, for eGFP - and Arch - expressing mice (black, and red traces; grey- and pink shades represent SEM). *Blue* windows indicate the 30-s tone blocks (CS), and *yellow* lines indicate overlapping footshocks and light stimulations. Horizontal *grey* lines denote the 30-s “no-CS” periods used for quantification in (F). (F) Quantification of freezing during the 30-s CS -, and no-CS epochs (*left* and *right* panels). No differences in freezing between eGFP and Arch - expressing mice was found (“n.s.”: not significant; see Results for statistical parameters). (G) Time-resolved freezing behaviour of eGFP- and Arch-expressing mice for the fear memory recall session. Note the robust CS - evoked freezing in the eGFP (control) mice, whereas the Arch - expressing mice show a conspicuous decrease in CS - evoked freezing (black, and red average traces, respectively). (H) Quantification of freezing during the CS (*left*) showed a significant impairment of freezing in the Arch - as compared to the eGFP - expressing mice, while freezing during the no-CS epochs (right) was unchanged (see Results for statistical parameters). (I) The CS - specific freezing, calculated as average freezing during the four CS epochs, *minus* the average freezing during the no-CS epoch, shows a significant decrease in the Arch mice as compared to the control mice (p = 0.0019, t-test). (D, I: bars indicate mean ± SEM; F, H: symbols indicate mean ± SEM)

To test the hypothesis that footshock-driven activity of principal neurons in the pInsCx contributes to the acquisition of an auditory-cued fear memory, we silenced these neurons precisely during the footshocks presented on the training day of a fear learning paradigm, by applying a 3s green light-pulse (561nm) starting 1s before each footshock (Figure 1C; see also Johansen et al., 2014). On the habituation day, the mice showed little freezing to the tone, and the amount of freezing was indistinguishable between the Arch- and the eGFP groups (Figure 1D; p=0.2373, U=109; Mann-Whitney U test). On the training day, when 3s green light-pulses (561nm) were applied during each footshock (Figure 1E; orange lines), mice in the Arch-, and the eGFP group showed a gradual increase in freezing during the training session. This acute freezing response was interrupted after each footshock, when the mice displayed increased locomotor activity and consequently, the freezing readout dropped; however, the acute freezing response did not appear to be different between mice in the Arch- or the eGFP group (Figure 1E). To analyze freezing quantitatively, we averaged the percent of the time mice spent freezing during 30-s “no-CS” intervals, and during the six CS periods (see Figure 1E, lower grey bars, and vertical blue bars, respectively). This did not reveal significant differences between the Arch- and the eGFP-expressing mice (Figure 1F; p=0.41, F(1, 32)=0.684 and p=0.84, F(1, 32)=0.04 for the CS- and no-CS periods respectively; two-way repeated measures ANOVA). Thus, footshock-driven activity of pInsCx principal neurons was not necessary for the freezing response of the mice during the training session.

One day later, the mice were presented with four CS stimuli in a different context, to test for recall of the auditory-cued fear memory. Control mice expressing eGFP showed a strong increase of freezing in response to each CS, indicating that they had formed a memory of the auditory cue (Figure 1G, black trace; N=16 eGFP-expressing mice). Arch-expressing mice, on the other hand, showed a weaker CS - induced freezing response (Figure 1G, red trace, N=17 Arch-expressing mice). Analysis during the 30-s CS and no-CS periods showed that freezing during the CS was significantly reduced in the Arch - expressing mice, as compared to the eGFP mice (Figure 1H, *left*; p=0.0094, F(1, 32)=7.64; two-way repeated measures ANOVA). Conversely, freezing during the no-CS epochs was not significantly different between the groups (Figure 1H; *right*, p=0.92, F(1, 32)=0.009, two-way repeated measures ANOVA). Similarly, the *CS - specific* freezing, defined as the difference in freezing between the CS and the no-CS epochs, was reduced in mice expressing Arch as compared to eGFP (Figure 1I, p=0.0022, t=3.33, df=32; t-test). These experiments show that footshock-evoked activity of pInsCx principal neurons is necessary for the acquisition of an auditory-cued fear memory.

### The pInsCx provides a major cortical input to the LA

We hypothesize that during fear learning, the pInsCx sends footshock information to downstream brain areas. Therefore, we next investigated the output connections of the pInsCx, using a virus-mediated anterograde labeling approach based on a Synaptophysin-mCherry construct (Materials and Methods). This approach revealed mCherry-positive nerve terminals in areas of the prefrontal cortex (infralimbic- and dorsal peduncular area; ILA and DP respectively; Figure 2B); in the Nucleus accumbens (ACB; Figure 2C); in the LA and in the adjacent tail striatum (LA and CP; Figure 2D); in the lateral BA (BLAp; Figure 2E); as well as in thalamic areas like the posterior thalamic nuclear group (PO), posterior thalamic nucleus, triangular part (PoT), the posterior intralaminar thalamic nucleus (PIL), and in the parvicellular part of the ventral posteromedial nucleus (VPMpc; see Figure 2F). Moreover, on the contralateral brain side, we found outputs from the pInsCx in the tail striatum, in the LA, in the posterior agranular insular cortex, and in the pInsCx (CP, LA, AIp and VISC; Figure S2A, B). On the other hand, the claustrum- and endopiriform nuclei adjacent to the LA were largely unlabeled (EPd and EPv; Figure 2D). Similar observations as in Figure 2B-F were made in N = 3 mice (see Table 1 for definitions of brain area terminology used here; Allen Brain Atlas [ABA], 2014).

**Figure 2.**
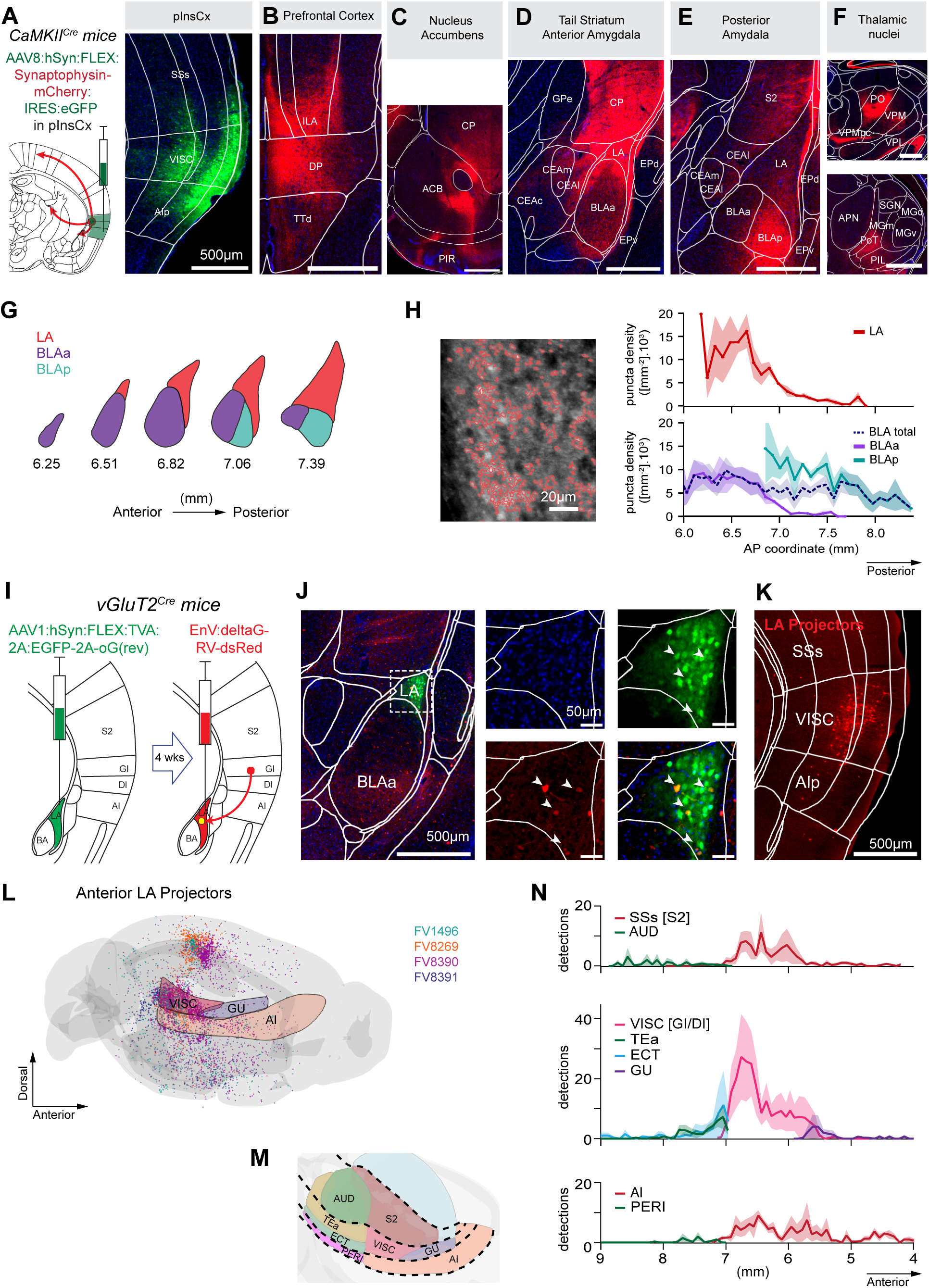
Virus-mediated antero- and retrograde labeling shows that the pInsCx provides a major input to the anterior LA. (A) *Left*, scheme of the experimental approach for anterograde labeling of outputs from the pInsCx. *Right*, eGFP fluorescence image from the center of the injection site. The images in (A-F), and in Figure S2I, J are from one representative mouse. (B - F) Example images showing Synaptophysin-mCherry fluorescence in the ventral prefrontal Cortex (B), in the Nucleus accumbens (C), in the tail striatum and anterior LA (D), in the more posteriorly located lateral BA (E), and in various thalamic nuclei (F). (G) Outlines of the ABA areas of the BLA complex, sub-divided in LA, BLAa, and BLAp along various a-p positions. (H) Analysis of the density of nerve terminals from the pInsCx in the three BLA substructures along the a-p axis. *Left*, example image of Synaptophysin-mCherry - positive nerve terminals in the LA, with superimposed detections (red lines). *Right*, Average nerve terminal density in the LA (*top*), and in the BLAa, BLAp, and the sum of BLAa and BLAp (“BLA total”; *bottom;* N = 3 mice). (I) Scheme of monosynaptically-restricted, rabies virus-mediated back-labeling of presynaptic neurons, using a VGluT2^Cre^ mouse to target excitatory neurons in the LA as starter cells (see also Figure S2A-D). (J) Images of eGFP - positive neurons expressing the helper construct (green channel), and dsRed - expression caused by a second injection of pseudotyped rabies virus four weeks later in the anterior LA (see also scheme in I). The images on the right show the anterior LA on an enhanced scale, with the DAPI channel alone (blue, top - left); the eGFP channel (green, top - right), the dsRed channel (red, bottom - left), and finally, the overlay of the dsRed and eGFP channel (bottom - right). A small number of eGFP - positive neurons in the anterior LA also expresses dsRed, thus defining the starter neuron population in the LA. (K) Example image of dsRed - positive, back-labeled neurons in the lateral cortex area. Note the cluster of LA - projectors in the pInsCx (called “VISC” in the ABA), whereas fewer dsRed - positive neurons are seen in the adjacent cortical areas. Brain section images were aligned to the Allen Brain Atlas (see Materials and Methods; see Table 1 for the meaning of abbreviations of brain structures). (L) 3D-rendering of all cortical neurons back-labelled from the anterior LA in N = 4 mice (mouse identity codes on the right). The volumes of the pInsCx GI and - DI (“VISC”), of the gustatory part of the insular cortex (“GU”), and of the agranular Insular cortex (“AI”) are also shown. (M) Outlines of lateral cortex brain areas according to the ABA, used for the analysis of a-p distribution of LA-projectors in (N). (N) Average number of detected neurons (N = 4 mice) in the lateral cortex along the a-p axis. Results are shown for the more dorsal SSs (S2) and AUD (*top*); for the VISC (pInsCx GI, - DI), TEa, ECT and GU (*middle*); and for the more ventral cortices of the AI and PERI (bottom). Note the peak of detected back-labeled neurons from the LA in the pInsCx - GI and - DI (middle panel, light red trace). (H, N: traces and lighter hues represent mean ± SEM. B - F: scale bars, 500 µm).

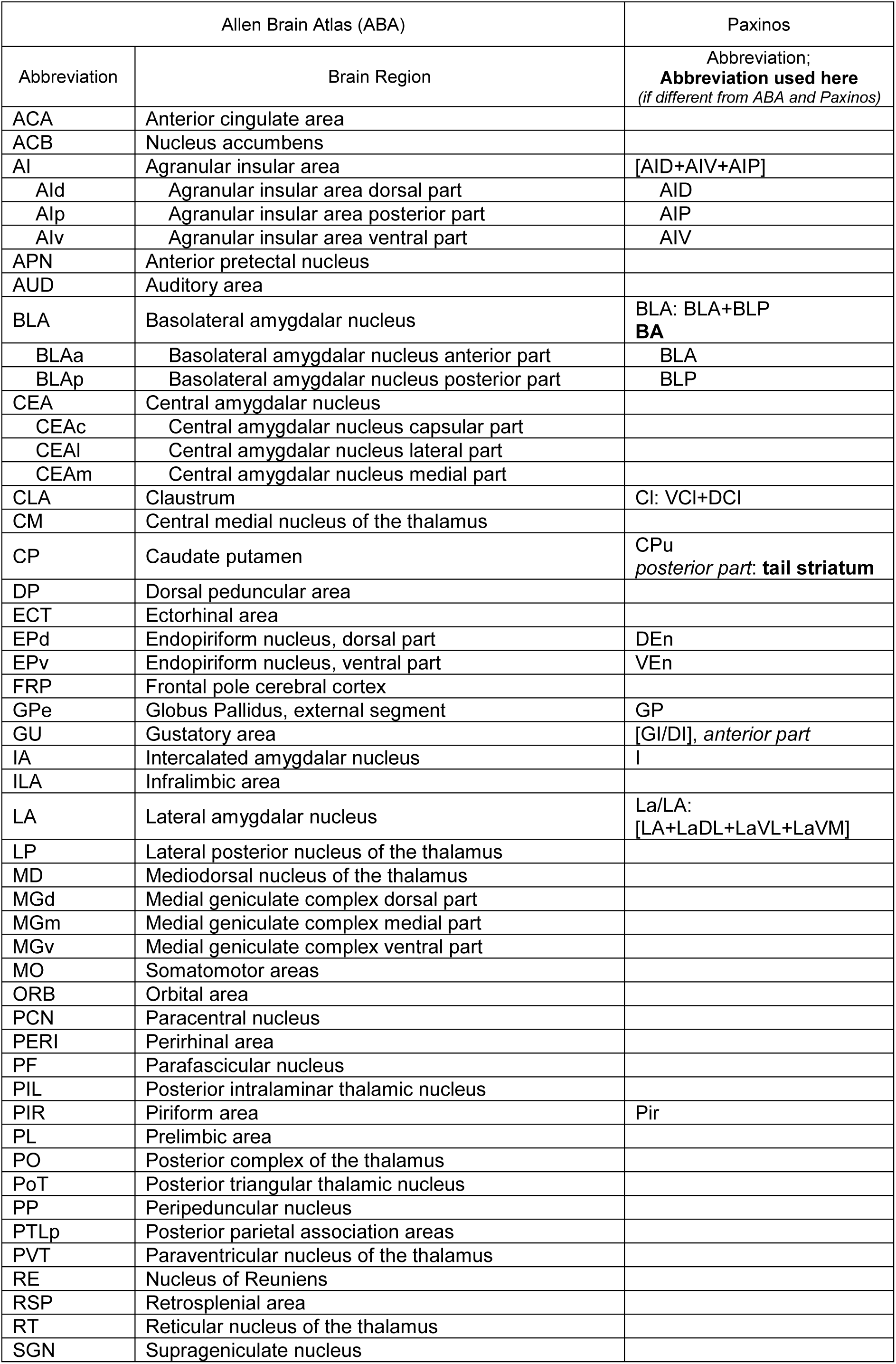
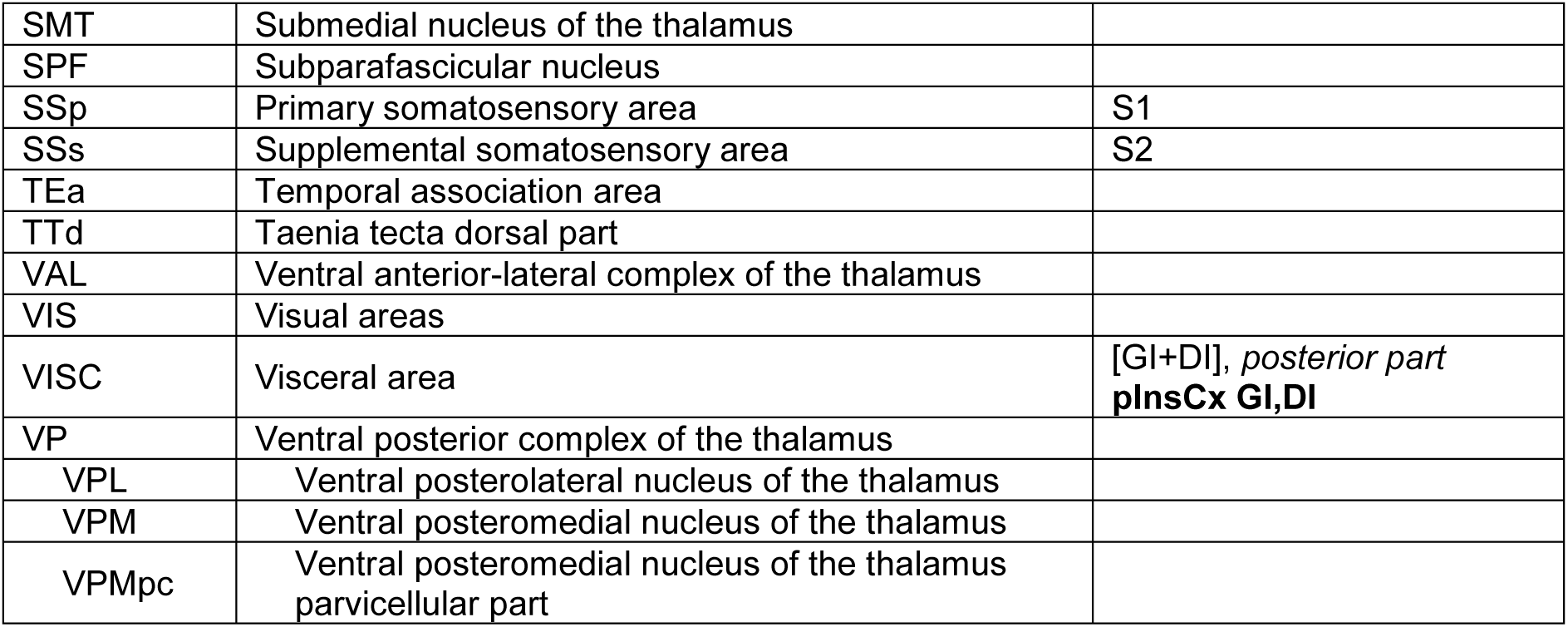
Table 1

Given the established role of the LA and BA in fear learning, we further investigated the outputs from the pInsCx to these basolateral amygdalar areas, analyzing the input synapse density along the a-p axis (Figure 2G, H). The nerve terminal density was high in the anterior LA, and dropped towards the more posterior LA (Figure 2H, red data). In the BA (divided into BLAa and BLAp in ABA terminology; see Table 1), nerve terminals from the pInsCx were observed in the BLAa, and more posteriorly in the BLAp; the latter corresponds to the lateral BA (Figure 2E; Figure 2H, blue average data trace). Thus, virus-mediated anterograde tracing shows that the pInsCx provides a prominent cortical input to the basolateral amygdala (BLA) complex, which mainly targets the anterior LA, and the lateral BA.

We next validated the anatomical finding of a pInsCx to LA projection, using rabies virus mediated, mono-synaptically restricted retrograde labeling (Wickersham et al., 2007). We employed VGluT2^Cre^ mice, to limit the expression of the helper virus to principal neurons of the LA, while excluding its expression in the neighboring posterior striatum (Figure S2A-D). Injections of helper virus into the anterior LA of VGluT2^Cre^ mice (AAV1:hSyn:FLEX:TVA:2A:eGFP:2A:oG) led to the expression of eGFP, and presumably TVA and oG (Figure 2J, green channel). Four weeks later, a pseudotyped rabies virus driving the expression of dsRed was injected into the anterior LA, leading to the co-expression of dsRed in a few eGFP-positive neurons, thereby defining the starter population of LA principal neurons (Figure 2J, arrows; Wickersham et al., 2007).

Analysis of the back-labelled neuronal populations, concentrating on thalamic- and cortical structures, revealed pools of back-labelled, dsRed - expressing neurons in various forebrain areas. In the thalamus, we observed the strongest back-labeling in the PO, in the mediodorsal nucleus (MD), in the ventral posteromedial nucleus (VPM) and in the PoT, suggesting that somatosensory-related thalamic nuclei provide input to the anterior LA (Figure S2E). In the cortex, the four areas with the largest number of back-labelled neurons were the primary somatosensory cortex (S1), the granular - and dysgranular part of the pInsCx (pInsCx - GI and - DI respectively, called VISC in the ABA), the agranular insular area (AI), and the secondary somatosensory cortex (S2; called SSs in the ABA; Figure 2K, L; Figure S2F). Because it is possible that the ABA reference atlas might not correctly capture the border between the pInsCx and the dorsally adjacent S2, we validated the alignments using two genetic mouse models expressing Cre-recombinase under the control of relevant marker genes (Figure S2G, H). These were Scnn1a^Cre^ x tdT mice which label cortical layer 4 neurons (Madisen et al., 2010; Vickers et al., 2018) - we found a cluster of Scnn1a+ neurons at the border between S2 and pInsCx (Figure S2G). Furthermore, we used Etv1^Cre^ x tdT mice which label layer 5 neurons (Yoneshima et al., 2006; Sürmeli et al., 2015), and found that a band of Etv1+ neurons expands in width in the pInsCx-GI and DI, correlating with the thinning of layer 4 in the pInsCx (Figure S2H). These findings confirm the assignment of the pInsCx (VISC) - S2 border in ABA brain atlas alignments.

Finally, we analyzed the number of back-labelled neurons along the a-p axis, concentrating on the lateral cortical areas (Figure 2M). In these areas, the highest number of LA - projectors were found in the pInsCx, followed by the S2 and by the AI (Figure 2N, red average data traces, N = 4 mice). On the other hand, the more posteriorly located ectorhinal cortex (ECT), temporal association cortex (TEa) and auditory cortex (AUD) contained fewer back-labelled neurons (Figure 7N, green average data traces, N=4 mice). Previous work has shown that the TEa and auditory cortices are major cortical inputs to the LA (Romanski and LeDoux, 1993; Dalmay et al., 2019). We attribute the different findings in the present study to the targeting of rabies virus to the anterior LA. Taken together, virus-mediated antero- and retrograde labeling techniques show that the pInsCx provides a major cortical input to the BLA complex, and there, mainly to the anterior LA and to the lateral BA (Figure 2).

### The pInsCx makes robust glutamatergic synapses in the LA

To characterize the pInsCx - LA connection functionally, we next employed optogenetically-assisted circuit mapping (Petreanu et al., 2007). We expressed the channelrhodopsin variant Chronos (Klapoetke et al., 2014) in principal neurons of the pInsCx on both brain sides, using CaMKII^Cre^ x tdT mice and a Cre-dependent expression vector (Figure 3A; Materials and Methods). Four weeks after surgery, we performed whole-cell recordings in slices of the LA, targeting principal neurons by means of their tdTomato - fluorescence (Figure 3B, right). Single blue light pulses (470 nm; ∼ 5 mW/mm^2^ intensity) caused optogenetically-evoked EPSCs (oEPSCs), with a fast component at negative holding potentials (−70 mV) and an additional slow component at + 50 mV (Figure 3C). The slow component was blocked by the NMDA-antagonist AP5 (D,L-2-amino-5-phosphonovaleric acid; 50 µM); the fast component measured at -70 mV was blocked by the AMPA-receptor antagonist NBQX (5 µM; n=3 cells; Figure S3A). These experiments identify the pInsCx - LA connection as a glutamatergic synapse.

**Figure 3.**
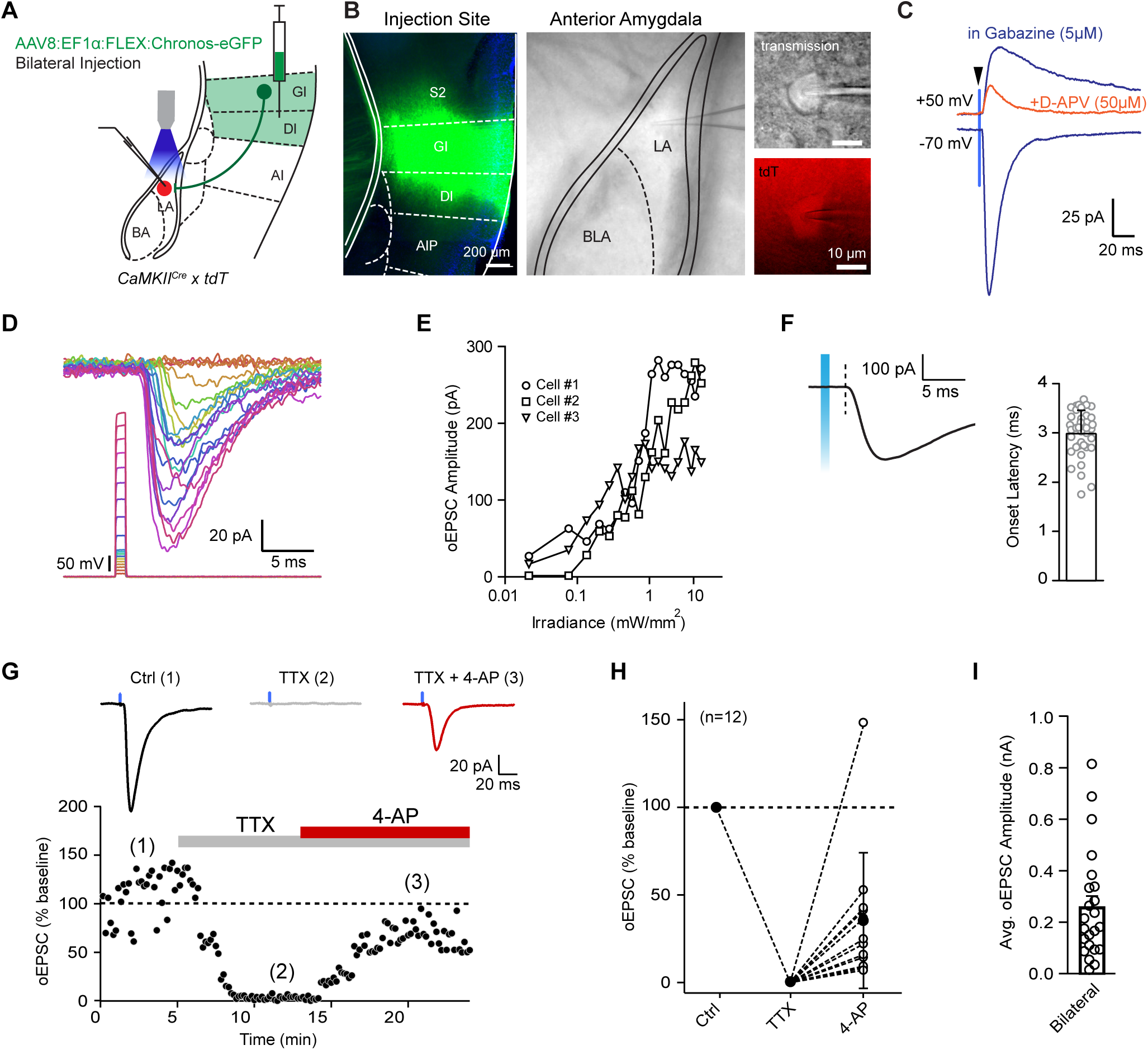
*Ex-vivo* optogenetic circuit mapping reveals a robust glutamatergic synapse from the pInsCx to the LA. (A) Scheme of the experiment design. (B) *Post-hoc* validation of Chronos-eGFP expression at the injection site in the pInsCx (*left*), and image of the recording site in the LA slice (*middle*), as well as zoom-ins of a recorded tdTomato positive LA principal neuron (*right*). (C) Blue light stimulation (1 ms; ∼5 mW/mm^2^; blue vertical line) evoked a fast-component EPSC at -70 mV, and a mixed fast - and slow EPSC as + 50 mV (blue traces; in the presence of 5 µM gabazine to block feedforward inhibition). Addition of 50 µM AP5 blocked the slow NMDA-EPSC component at +50 mV (red trace). (D) Gradually increasing the stimulus light intensity led to a gradual increase of oEPSC amplitudes. (E) Input-output relationship of oEPSC as a function of light intensity (n = 3 example recordings). (F) Illustration of the oEPSC latency (*left*), and average and individual data points (*right*; n = 41 recordings from N = 10 mice). (G) Time plot of oEPSC amplitudes recorded in an example principal LA neuron upon 1 ms light stimulation under control conditions (1), after application of 1 µM TTX (2), and after co-application of a 4-AP (1 mM) in the continued presence of TTX. (H) Quantification of average oEPSC amplitudes under the three conditions (n = 12 recordings from N = 5 mice). Note the partial recovery of oEPSCs after co-application of 4-AP. (I) Average and individual data points for oEPSC amplitudes recorded after bilateral, Cre-dependent expression of Chronos in CamKII^Cre^ mice (see panel A; n = 23 recordings from N = 6 mice). (F, I: bars indicate mean ± SEM)

We next measured input - output relations by changing the intensity of the stimulus light, to study the convergence of presynaptic axons onto LA principal neuron (see also Litvina and Chen, 2017; Gjoni et al., 2018). Increasing the light intensity gradually increased the oEPSC amplitudes without discernible steps, indicating that the oEPSCs were composed of several unitary EPSCs with smaller amplitudes (Figure 3D, E). The oEPSCs had a fast onset latency (3.11±0.07 ms, n=41 recordings) and a smooth rising phase (Figure 3F). Furthermore, the oEPSCs were blocked by tetrodotoxin (TTX, 1 µM), and this block was partially recovered when the K^+^-channel blocker 4-aminopyridine (4-AP, 1mM) was co-applied in the continued presence of TTX (n=12 recordings; Figure 3G, H). These findings suggest that the pInsCx makes a direct, monosynaptic connection onto LA principal neurons.

The average oEPSC amplitude upon near - maximal light intensity was 257 ± 44 pA (Figure 3I; n = 23 recordings from N = 6 mice with bilateral, Cre-dependent expression of Chronos). Additional experiments showed that oEPSCs of nearly identical amplitude were obtained when Chronos was expressed Cre-independently; that unilateral expression of Chronos led to significantly smaller oEPSCs; and that a larger connectivity and oEPSC amplitudes were found in more anterior - as compared to more posterior slices of the LA (Figures S3B-D). These experiments agree with our anatomical observations (Figure 2), and establish that the pInsCx makes a robust glutamatergic, excitatory connection to the LA.

### The pInsCx - LA synapse undergoes plasticity upon fear learning

We next investigated whether the glutamatergic connection from the pInsCx - LA undergoes long-term plasticity after auditory-cued fear learning. Measurements of the paired-pulse ratio, and of the AMPA/NMDA ratio following fear learning, can be used to assess whether learning induces long-term plasticity at a given synaptic connection (Lucas et al., 2016; Yu et al., 2017; Rich et al., 2019). For this, we injected AAV8:hSyn:Chronos bilaterally into the pInsCx of CaMKII^Cre^ mice; tdTomato was additionally expressed Cre-dependently in the LA to target principal neurons in the subsequent recordings (Figure 4A, B). Three weeks later, mice were randomly assigned to two groups: a control group in which mice were exposed only to the CS (“CS-only” group), and a group that underwent the standard fear learning protocol (“CS+US”; Figure 4C). Mice in the CS-only group showed negligible freezing both during the training- and the recall sessions, whereas mice in the CS+US group showed CS-induced freezing during the recall session, which suggests that they have formed an auditory-cued fear memory (Figure S4).

**Figure 4.**
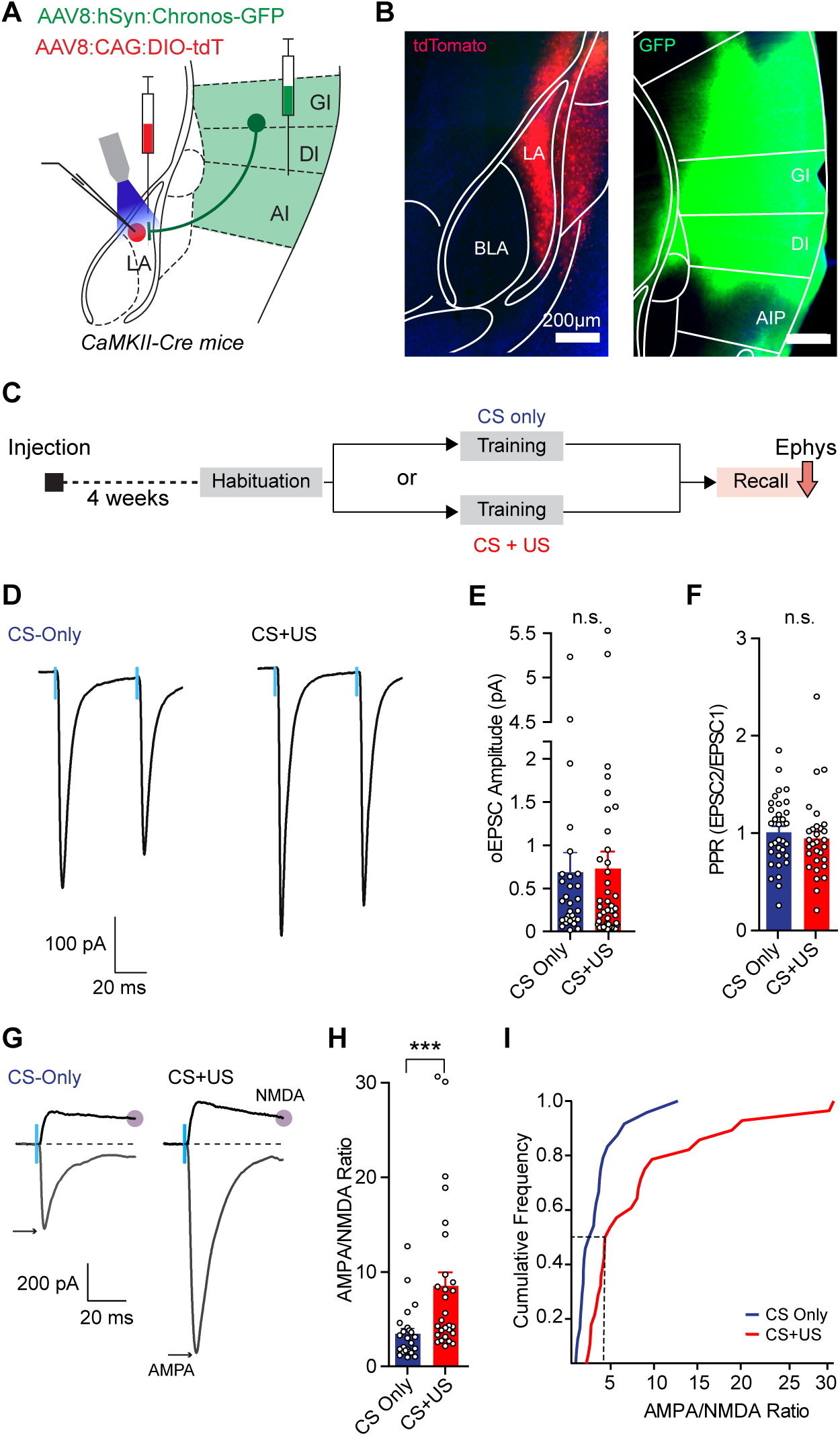
Postsynaptic long-term plasticity induced by auditory-cued fear learning at the pInsCx - LA synapse. (A) Scheme of the bilateral expression of Chronos in the pInsCx for the later recording of oEPSCs in the LA. (B) Example fluorescence images to validate injection targeting in the LA (left) and in the pInsCx (right). (C) Scheme of assigning two behavioral groups. (D) Example oEPSCs traces recorded in the LA upon paired stimulation (interval, 50 ms), in the two experimental groups. (E, F) Average and individual data points for the oEPSC amplitudes (E) and paired-pulse ratios (F) in the two experimental groups. Neither of the measures were significantly different between the groups. (G) Representative oEPSC traces for each group, recorded at -70mV to measure the AMPA-component, and at +50mV to measure the NMDA-component (see grey circle, 50ms after light onset). (H, I) Average and individual data points (H), and cumulative distributions of AMPA/NMDA ratios (I). Note the significant increase in AMPA/NMDA ratio in the CS+US group, as compared to the CS-Only group (p = 0.0002, Mann Whitney U test). (E, F, H: bars indicate mean ± SEM. D, G: average traces from n = 10 subsequent traces).

Following the fear memory recall session, the mice were sacrificed, and oEPCSs were measured at the pInsCx - LA connection. We found that the oEPSC amplitudes, and the PPR were indistinguishable between the two groups (Figure 4D-F; p = 0.89, U=573 and p=0.239, U=409 respectively, Mann Whitney U test). On the other hand, the AMPA/NMDA ratio was significantly larger for oEPSCs recorded in the CS+US group, as compared to the CS-only group (Figure 4G, H; p=0.0002, U=142, Mann Whitney U test). Cumulative distributions of the AMPA/NMDA ratios show a broader distribution, and the occurrence of large AMPA/NMDA ratios in a subset of the investigated LA principal neurons in the CS+US group (Figure 4I). Thus, during auditory-cued fear learning, the pInsCx - LA connection undergoes a postsynaptic form of LTP.

### Footshock - driven activity of LA-projectors is necessary for acute freezing and for efficient fear learning

We showed that footshock-driven activity of principal neurons in the pInsCx is necessary for the efficient formation of a fear memory (Figure 1), and that the pInsCx makes a robust glutamatergic connection to the LA (Figures 2, 3). We therefore asked whether, more specifically, the activity of LA - projectors during footshocks is necessary for the formation of an auditory-cued fear memory. For this, we used a retrograde expression approach, in which AAVretro:EF1α:mCherry:IRES:Cre was injected bi-laterally into the LA, combined with the Cre-dependent, bilateral expression of Arch in the pInsCx (AAV1:CAG:FLEX:Arch-eGFP; Figure 5A). As before, mice in a control group expressed eGFP in a Cre-dependent manner bilaterally in the pInsCx, but underwent otherwise the same procedures as the Arch - expressing mice. An unavoidable spill-over of the AAVretro vector into the claustrum / endopiriform area lateral to the LA (Figure S5) should not affect the conclusions of the behavioural experiments, because pInsCx axons spare this area (Figure 2D). On the other hand, our targeting approach of the AAVretro vector injection avoided the posterior striatum medial to the LA (Figure S5), making it unlikely that posterior striatum projectors in the pInsCx contribute to the observed behavioral effects.

**Figure 5.**
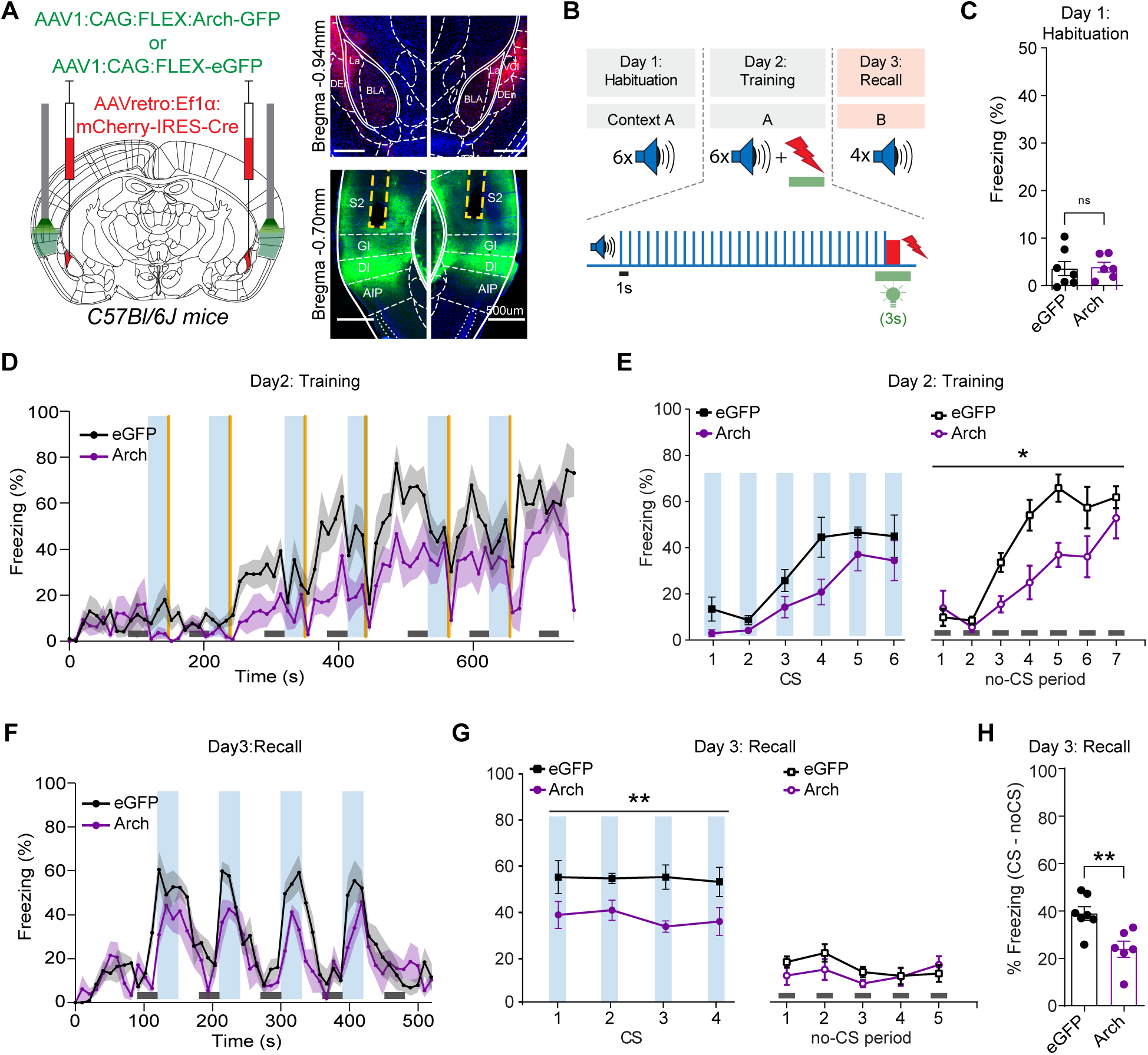
Optogenetic silencing of LA-projectors re-capitulates the fear memory deficit observed after silencing pInsCx principal neurons. (A) *Left*, scheme of the bilateral retrograde expression approach of Arch-eGFP in LA-projectors of the pInsCx, and fiber positioning bilaterally in the pInsCx. *Right*, example images of mCherry expression in the LA (*top*), and Arch-eGFP expression in the pInsCx (*bottom*). (B) Scheme of the timing of Arch - mediated inhibition during the footshocks on the training day. (C) Freezing during the CS blocks on the habituation day for Arch - and eGFP - expressing mice (N = 6 and 7; purple and black data points; p = 0.877, t-test). (D) Time-resolved average freezing behavior for N = 6 Arch - and N = 7 control mice during the training session (purple, and black traces). *Blue* windows indicate the CS; *yellow* bars indicate the time of green light application during the footshocks. (E) Quantification of freezing during the 30-s CS - and no-CS epochs (*left* and *right* panel, respectively). During the no-CS periods, freezing was significantly lower in the Arch group as compared to the eGFP group (p = 0.008, 2-way repeated measures ANOVA; see Results for further statistical parameters). (F) Time-resolved freezing of eGFP- and Arch mice during fear memory recall. (G) Quantification of freezing during the 30-s CS blocks (*left*), and during the 30-s no-CS periods (*right*) during fear memory recall. Freezing during the CS was significantly smaller in the Arch group as compared to the eGFP group (see Results for statistical parameters). (H) Average and individual data for CS - specific freezing. Arch mice had a significantly decreased CS - induced freezing as compared to the control mice (p = 0.006, t-test). (C, H: bars indicate mean ± SEM. D, F: data points and light color hues indicate mean ± SEM. E, G: data points and bars indicate mean ± SEM).

In order to silence the US - driven activity of LA-projectors, we again applied green light for 3s, starting 1s before each footshock presentation on the fear learning training day (Figure 5B). On the habituation day, the mice showed little freezing, and the freezing was indistinguishable between the groups (Figure 5C; p=0.877, t=0.1584, df=11, t-test). On the training day, the freezing response of the mice in the Arch group appeared weaker than in the control mice (Figure 5D, violet- and black trace; N=6 and 7 mice in the Arch- and the eGFP group, respectively). Time-averaging over 30-s time bins showed that during the no-CS periods, mice in the Arch group froze significantly less than mice in the control group (p=0.008, F(1, 11)=10.4; two-way repeated-measures ANOVA; Figure 5E, *right*). During the CS periods, freezing in the Arch group was also smaller than in the control mice, but this difference did not reach statistical significance (Figure 5E left; p=0.06, F(1, 11)=4.33; two-way repeated-measures ANOVA). Thus, footshock - driven activity of LA - projector neurons in the pInsCx contributes to the freezing response of mice during the fear learning training day.

During the fear memory recall session, Arch mice showed a reduced amount of freezing in response to the CS presentations as compared to control mice (Figure 5F). Time-averaged quantification revealed that freezing during the CS was significantly reduced in the Arch group (Figure 5G, *left*; p=0.0017, F(1, 11)=17.09; two-way repeated measures ANOVA; N=6 and 7 mice in the Arch - and the eGFP group, respectively). On the other hand, freezing during the no-CS epochs was low in both experimental groups, and not significantly different (Figure 5G, *right*; p=0.44, F(1, 11)=0.636; two-way repeated measures ANOVA). Furthermore, the CS - specific freezing was significantly reduced in the Arch group as compared to the control group (Figure 5H; p=0.006, t=3.391, df=11; unpaired t-test). Taken together, these experiments expand the findings of Figure 1, and show that footshock-driven activity of a sub-population of the pInsCx principal neurons, i.e. the ones projecting to the LA, is necessary for the efficient formation of an auditory-cued fear memory.

### pInsCx drives footshock responses in a subset of LA neurons

It has been shown that sub-populations of LA neurons respond to footshock stimulation (US; Gore et al., 2015; Grewe et al., 2017), and that footshock-induced depolarization of LA neurons contributes to auditory-cued fear learning (Johansen et al., 2014). We found that neurons in the pInsCx make a robust glutamatergic synapse onto LA principal neurons, and that the footshock-driven activity of these LA-projectors contributes to fear learning (Figures 3, 5). Therefore, we hypothesize that pInsCx neurons contribute to driving footshock responses in the LA. To investigate this possibility, we performed *in-vivo* Ca^2+^ imaging of the activity of LA neurons during fear learning, combined with optogenetic inhibition of pInsCx axons in the LA.

We used a dual wavelength miniature microscope, which enables simultaneous activation of a Ca^2+^ indicator and an inhibitory opsin, using light of 450 nm and 630 nm excitation wavelength, respectively (Stamatakis et al., 2018; Materials and Methods). We imaged neurons in the anterior LA, which, as we showed above, receives a dense innervation from the pInsCx (Figure 2, 3). We Cre-dependently expressed a genetically-encoded Ca^2+^ indicator in the left anterior LA of CaMKII^Cre^ mice. Furthermore, a virus driving the expression of Halorhodopsin (AAV8:hSyn:eNpHR3.0-eYFP) was injected bi-laterally in the pInsCx, allowing the expression and transport of Halorhodopsin in pInsCx axons innervating the LA. Finally, a GRIN lens was implanted over the left LA in the same surgery (Figure 6A). Four weeks later, LA neurons were imaged during the three-day fear learning protocol. A 3s pulse of orange light (wavelength 630 nm) was applied during the first, third, and fifth footshock (US) to activate Halorhodopsin, whereas the other US presentations were used as internal controls (Figure 6B).

**Figure 6.**
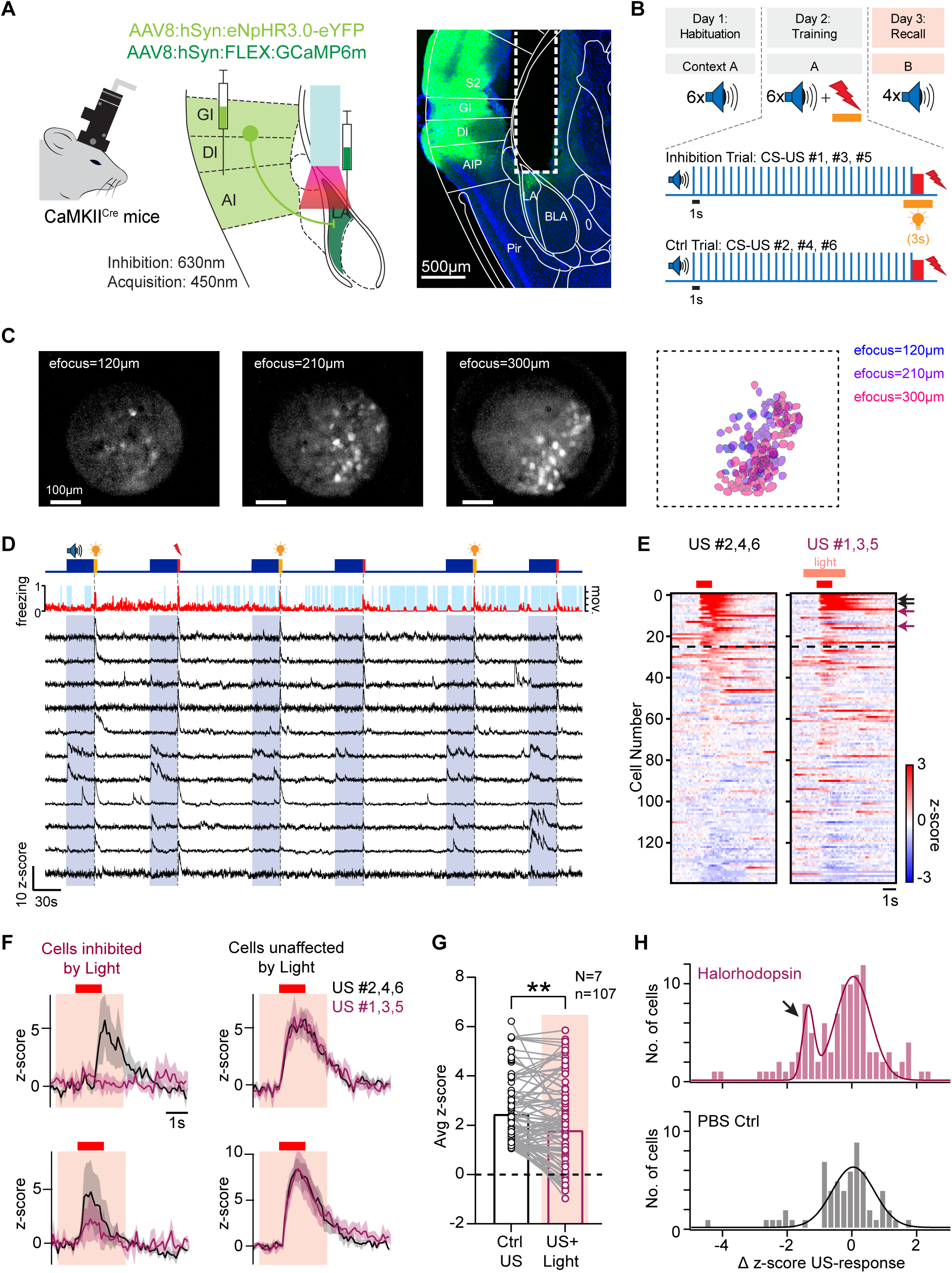
*In-vivo* Ca^2+^ imaging and optogenetic axonal silencing shows that the pInsCx drives footshock responses in a sub-population of LA neurons. (A) *Left*, scheme of *in-vivo* Ca^2+^ imaging in the LA with GCaMP6m, and simultaneous optogenetic inhibition of input axons from the pInsCx (with Halorhodopsin, “eNpHR3.0”). *Right*, Post-hoc histological validation of GCaMP6m and Halorhodopsin-eYFP expression, and placement of the GRIN lens over the anterior LA. (B) Experimental strategy to combine Ca^2+^ imaging and optogenetic silencing during the first, third, and fifth footshock application of the training session. (C) *Left*, maximal intensity projection images of GCaMP6m fluorescence at three focal planes used here (see Materials and Methods). *Right*, overlay of the imaged neurons extracted from the three focal planes. (D) The movement trace of an example mouse (*red*) and the freezing times (*light blue*; *top*) during the training session, as well as example Ca^2+^ traces from n=11 imaged LA neurons (*black; bottom*). Application of CS blocks (blue), footshocks (red, after each CS), and orange light (orange) is indicated in the top panel. (E) Color - coded plot of z-scored Ca^2+^ traces for all n=139 cells imaged in an example mouse, for US #2, 4, 6 (no light; *left*), and for US #1, 3, 5 (with orange light to activate Halorhodopsin; *right*). Data were sorted according to their average US-response to US#2, #4 and #6. Cells above the dashed line were classified as US-responders. Note cells with a clear suppression of the footshock response (purple arrows), whereas the Ca^2+^ transients in other cells were unaffected (black arrows). (F) Example Ca^2+^ responses of the cells indicated by arrows in (E). Two cells showed a decreased Ca^2+^ transient upon activation of Halorhodopsin (*left* panels), whereas another two example cells showed no change in Ca^2+^ (*right*). (G) Footshock-induced Ca^2+^ transient amplitude in n = 107 US - responsive cells imaged in N = 7 mice, averaged for footshocks #2, 4, 6 (no light; black data points, *left*), and for footshocks #1, 3, 5 (with orange light; purple data points, *right*). There was a significant decrease of the footshock - driven Ca^2+^ transient amplitude between the two conditions (p = 0.0056, Wilcoxon test). (H) Plots of Δ z-score^US^ (see Results for definition), indicating the degree of modulation of the Ca^2+^ transient by light application. Note the distinct peak of neurons with a negative Δ z-score^US^ response, observed in N = 7 mice in which Halorhodopsin was expressed in the pInsCx (top panel; data required a double-Gaussian for fitting). No second peak was observed in a PBS-injected control group (N = 4 mice, *lower* panel, grey data, requiring a single Gaussian for fitting). See also Figure S6. (data in A, and C - F are from the same example mouse).

During the fear learning training day, LA neurons showed varying activity patterns, with footshock responses in a sub-population of the imaged neurons, and Ca^2+^ responses during the CS in some neurons (Figure 6D; Grewe et al., 2017; Zhang and Li, 2018). We found that 107 out of 448 imaged neurons, or 23.8%, displayed detectable footshock responses (N = 7 Halorhodopsin - expressing mice). A plot of z-scored Ca^2+^ traces for all imaged LA neurons in one example mouse suggests that some neurons had reduced US-responses upon optogenetic inhibition, whereas other neurons were un-affected by light (Figure 6E, violet - and black arrows respectively; see Figure 6F for the corresponding example traces). Averaging the Ca^2+^ amplitudes in all imaged neurons revealed a significant reduction of the amplitude of footshock - driven Ca^2+^ signals (Figure 6G; p=0.0056, W=-1772; Wilcoxon test; n=107 US-responsive neurons in N=7 mice). Furthermore, calculating the difference between the Ca^2+^ transient amplitude in the presence *minus* the absence of light (Δ z-score^US^ = z-score^US, light^ - z-score^US, no light^), showed that the modulation of the Ca^2+^ transient amplitude scattered around zero in many neurons, but that in about one-fifth of the US-responsive neurons, the Δ z-score^US^ values were lower, thus producing a distinct, second peak in the histogram (Figure 6H *top*, arrow). To exclude a direct effect of orange light on the Ca^2+^ transient amplitudes, we performed control experiments in which mice received a saline injection into the pInsCx instead of the Halorhodopsin vector, and were then imaged in the same way as the mice expressing Halorhodopsin. In the control mice, application of orange light did not lead to a significant reduction of the Ca^2+^ transient amplitude in LA neurons (n = 73 US - responsive neurons in N=4 PBS - injected mice; p=0.07, W=-235; Wilcoxon’s test, Figure S6E). Furthermore, the Δ z-score^US^ values were evenly distributed around the origin, but did not show a second peak at lower values, as observed in the mice expressing Halorhodopsin (Figure 6H). Taken together, *in-vivo* Ca^2+^ imaging combined with optogenetic nerve terminal inhibition shows that the pInsCx - LA synapse drives US - responses in a sub-population of LA neurons during the fear learning training session.

## Discussion

The posterior insular cortex (pInsCx) processes somatosensory, nociceptive and temperature information (Rodgers et al., 2008; Benison et al., 2011; Vestergaard et al., 2022), and human imaging studies showed that the pInsCx is part of a larger network that processes nociceptive information (Apkarian et al., 2005; Coghill, 2020; Kuner and Kuner, 2021). Nevertheless, no clear evidence for a role of this brain area in fear learning has been obtained (Shi and Davis, 1999; Brunzell and Kim, 2001; Lanuza et al., 2004; Gehrlach et al., 2019). We have used tools for circuit investigation to study the role of the pInsCx in fear learning, and to identify relevant output pathways for this function. We show that footshock-driven activity of pInsCx principal neurons is necessary for the efficient formation of an auditory-cued fear memory. We then found that the pInsCx projects to specific components of the BLA complex and there, at high density to the anterior LA, besides projecting to other specific forebrain areas. The connection to the anterior LA provides robust, glutamatergic EPSCs to LA principal neurons, and the EPSCs at this connection undergo LTP during fear learning. Combined optogenetic axon silencing and *in-vivo* Ca^2+^ imaging showed that the pInsCx transmits US information to a subset of LA principal neurons. Finally, suppressing the footshock-evoked activity in LA-projectors recapitulates the deficit in fear memory formation found after silencing pInsCx principal neurons. Thus, our study suggests that the pInsCx processes behaviorally relevant information about acute nociceptive and aversive somatosensory events, and transmits this information via a plastic synapse to the LA. This identifies one source of US-input to LA neurons during auditory-cued fear learning.

### Suppression of footshock coding reveals mechanisms of fear memory acquisition

We used optogenetic suppression of neuronal activity during the footshock presentation, to investigate whether the pInsCx processes footshock information relevant for fear learning. This caused a specific deficit in CS - induced freezing during fear memory recall one day later (Figure 1). In control experiments, activation of Arch led to a significant suppression of footshock-driven activity of pInsCx neurons *in-vivo*. These experiments strongly suggest that excitatory neurons in the pInsCx process footshock information relevant for the acquisition of a fear memory.

A previous study in rats inactivated the pInsCx by muscimol before the training session, and observed a decreased fear memory recall (Casanova et al., 2016), which similarly indicates a role in fear memory acquisition. The previous study also made post-training manipulations, and concluded overall that the pInsCx is involved in the *consolidation* of a fear memory (Casanova et al., 2016); thus the pInsCx might have a role both in the acquisition, and in the consolidation of an auditory-cued fear memory. A recent study in mice showed that the pInsCx codes for longer-lasting aversive states; however, prolonged optogenetic inhibition did not reveal a role of this brain area in fear learning (Gehrlach et al., 2019). We see two possible reasons for this apparent discrepancy. First, differences in the optogenetic silencing protocols might be responsible. Gehrlach et al. (2019) used prolonged continuous light application (∼ 20 min) which might lead to partial inactivation of the inhibitory opsins; furthermore, it was not reported whether the footshock-driven activity of pInsCx neurons was suppressed by this manipulation; the present- and previous study also used different optogenetic tools (Arch, versus Halorhodopsin). Second, differences in the targeted sub-areas of the pInsCx seem likely. We investigate a somatosensory - auditory region in the pInsCx-GI / DI, that partially extends into the ventral S2 (Figures 1, S1). This area contains neurons coding for footshocks, movement-onset, and tones (D. Osypenko, S.P., O.K. and R.S., *unpublished*), as well as neurons that project to the LA (Figure 2L-N); furthermore, this area might overlap with a pInsCx area containing temperature-coding neurons (Bokiniec et al., 2022; Vestergaard et al., 2022). Conversely, the pInsCx investigated by Gehrlach et al. (2019) contains neurons that project to the CeA but only few neurons that project to the LA, and seems to be located more ventrally. Fitting with this view, CeA projectors of the pInsCx are located more ventrally than LA - projectors (*unpublished observations*). Taken together, we do not think that the two studies strongly contradict each other. Gehrlach et al. (2019) emphasized a prolonged coding of aversive states in the pInsCx; the present study shows that coding of acute nociceptive signals by a likely more dorsal pInsCx area contributes to the acquisition of a fear memory.

### Posterior insular cortex provides a robust excitatory synapse to the anterior LA

Because we found a role of the pInsCx in fear memory acquisition, we next determined which forebrain structures are targeted by the insular cortical area that we have studied. This revealed, in general, characteristic forebrain areas targeted by the pInsCx. These include the ventral prefrontal cortex (Yasui et al., 1991), nucleus accumbens (Gehrlach et al., 2019; Wang et al., 2022), anterior insular cortex (data not shown; Kimura et al., 2010), and thalamic nuclei with various sensory functions like the PO (Bokiniec et al., 2022), PoT (Gauriau and Bernard, 2004), VPMpc (Chen et al., 2011; Samuelsen et al., 2013), and MGm/PIL (Figure 2F), thus reflecting the multisensory nature of the pInsCx (Gogolla, 2017). Furthermore, these output areas confirm that we target the pInsCx in our study. In addition, we found a dense output to the tail striatum (TS) and to the neighboring anterior LA (Bokiniec et al., 2022), as well as outputs to the lateral BA, and weaker outputs to the CeA (Figure 2D, E). Interestingly, neurons of the direct-and indirect pathway in the ventral TS were recently found to receive strong glutamatergic input from the pInsCx, and to differentially regulate the expression of fear memory in the presence, and absence of a learned CS (Kintscher et al., 2023).

Given the known role of the LA in fear learning and the integration of US- and CS signaling, we further studied the outputs to the LA. To confirm the presence of LA-projectors in the pInsCx, we used rabies-virus mediated back-labeling and limited the starter cells to glutamatergic (VGluT2+) LA neurons, thus excluding neighboring vTS neurons which are GABAergic (Valjent and Gangarossa, 2021; Figure S2A). This revealed a high number of back-labelled neurons in the pInsCx and other somatosensory cortical and thalamic areas, suggesting that somatosensory-related areas preferentially connect to the anterior LA (Figure 2).

Recent anatomical input - output tracing of the mouse insular cortex found only a weak, and somewhat variable output from the pInsCx to the LA, whereas outputs to the CeA were clearly observed (Gehrlach et al., 2019; Gehrlach et al., 2020; see also Schiff et al., 2018; Ponserre et al., 2020 for functional outputs to the CeA). The fact that Gehrlach et al. (2019, 2020) did not observe a strong projection from the pInsCx to the LA is consistent with the assumption that we have targeted a more dorsal area of the pInsCx as compared to their studies (see discussion above). Finally, we used optogenetically assisted circuit mapping, which revealed robust functional glutamatergic synapses established by the pInsCx onto LA principal neurons, with a high connectivity especially in the anterior LA (Figures 3, S3). Thus, anatomical-and functional mapping unambiguously establish an excitatory projection from the pInsCx to the LA.

### Transmission of US-information at a plastic insula - LA synapse

Given that US-driven activity of pInsCx principal neurons contributes to fear memory acquisition, and that this cortical area makes a robust excitatory connection to the LA, we hypothesized that footshock information is transmitted from the pInsCx to the LA. To test this idea, we employed *in-vivo* Ca^2+^ imaging combined with optogenetic silencing of pInsCx axons. We found that about 25% of the imaged LA neurons showed footshock responses, similar to recent *in-vivo* imaging studies performed more ventrally at the LA - BA border (Grewe et al., 2017; Zhang and Li, 2018). Axon silencing led to a distinct down-regulation of the Ca^2+^ transients in about one-fifth of the US-responsive neurons (Figure 6). Halorhodopsin has been used in some studies to silence axonal inputs (Barsy et al., 2020), but its efficiency for presynaptic silencing has not often been tested (Mahn et al., 2016). In our data, a downregulation of the Ca^2+^ transient was more often observed in neurons showing moderate-rather than high-amplitude Ca^2+^ transients (Figure 6E, top), which suggests that saturation of the Ca^2+^ indicator might limit the detection of a modulation of the Ca^2+^ signal. Both factors indicate that the percentage of US-responsive neurons driven by the pInsCx inputs might be a lower-limit estimate. On the other hand, the persistence of Ca^2+^ transients in many US-responsive neurons upon optogenetic inhibition of pInsCx axons clearly suggests that other afferent synapses carry footshock information to the LA, like the input from the non-lemniscal auditory thalamus (Barsy et al., 2020), or other, not yet identified inputs.

Is the transmission of US-information from the pInsCx to the LA behaviorally relevant for learning? We tested this by soma silencing of the footshock response in LA-projectors of the pInsCx (Figure 5). This re-capitulated the deficits in fear learning seen after footshock silencing of pInsCx neurons (Figure 1), showing that LA-projectors are involved in the acquisition of a fear memory, likely by transmitting US - information to a sub-population of LA neurons. Silencing the LA - projectors during the footshock led, in addition, to reduced freezing on the training day. Nevertheless, the relevance of the acute freezing response on the training day for the learning process, and therefore for the development of a conditioned response to the CS one day later, remains unknown (Fanselow, 2018). A limitation in the experimental approach of retrograde expression of Arch is a potential spill-over to neighboring brain areas. Furthermore, if LA-projectors make axon collaterals in other brain areas, footshock-driven activity of LA-projectors will also influence US-signaling in those brain areas. Despite these unknown factors, the finding that LA - projector silencing causes a similar impairment of auditory - cued fear memory as more widespread silencing of pInsCx principal neurons, strongly suggests that footshock information is transmitted to the LA, where it helps forming an auditory-cued fear memory.

We also observed that fear learning causes an increase in the AMPA/NMDA ratio at the pInsCx - LA synapse, suggesting a learning-induced postsynaptic form of LTP (Figure 4). Signs of postsynaptic plasticity after fear learning have been found originally at the thalamus - LA connection (Rogan and LeDoux, 1995; McKernan and Shinnick-Gallagher, 1997; Rumpel et al., 2005; Nabavi et al., 2014) and at the auditory cortex to LA connection (Nabavi et al., 2014; Kim and Cho, 2017); however, signs of LTP were also found at the output synapses of the LA in the CeA (Li et al., 2013; Hartley et al., 2019). Thus, there is a larger number of long-range connections in the limbic system which undergo long-term plasticity after fear learning than originally thought (see also Palchaudhuri et al., 2022 for review). We can currently only speculate about the role of LTP at the pInsCx - LA synapse. On the one hand, LTP at this synapse might contribute to the formation of an auditory-cued fear memory. In this case, CS - information should be transmitted at this connection as well. Indeed, it has been shown that the pInsCx contains a subarea sensitive to auditory stimulation (Rodgers et al., 2008; Sawatari et al., 2011; Gogolla et al., 2014); these studies, in the light of our findings might prompt further investigations into CS-signaling in the pInsCx. On the other hand, it remains possible that the induction of LTP at this connection serves to enhance the transmission of US-information from the pInsCx to the LA upon repeated stressful experiences. Altogether, the finding that the pInsCx - LA connection shows plasticity after fear learning confirms an involvement of this connection in fear learning.

### Outlook and conclusions

Recent studies in mice have found a role of the insular cortex in homeostatic feedback regarding physiological need states, or bodily reactions to fear states. Thus, the insular cortex was activated by hunger-driven signals originating from the hypothalamus (Livneh et al., 2017), and received feedback from the vagus nerve about bodily expressions of fear; this feedback was found to influence the degree of fear memory extinction (Klein et al., 2021). Furthermore, classical *in-vivo* recordings in rats coupled with measurements of physiological parameters showed that the pInsCx both codes for changes of visceral- and cardiovascular parameters (Cechetto and Saper, 1987); and in turn can exert a top-down influence on cardiovascular function (Yasui et al., 1991). We find that the pInsCx signals acute nociceptive information (Rodgers et al., 2008) to the LA, to contribute to the acquisition of a fear memory. Given the recent finding that a partially overlapping pInsCx area is the cortical site coding for non-noxious skin temperature changes (Vestergaard et al., 2022), it will also be interesting to investigate whether the pInsCx is involved in processing noxious heat, and whether it can transmit such information to the amygdala as a teaching signal for learning-induced plasticity. Taken together, the insular cortex detects signals both from visceral sites (see references above), as well as somatosensory, nociceptive and thermal information from the body surface and thus, exteroceptive signals (Rodgers et al., 2008; Vestergaard et al., 2022); the latter can be used as a teaching signal to induce learning and plasticity in the BLA (this study). Given that the activity of the pInsCx is altered in multiple forms of psychiatric disease (Shepherd et al., 2012; Di Martino et al., 2014; Uddin, 2015), it will be important to further investigate the circuit organization of the pInsCx, to better understand how the pInsCx surveys both intero- and exteroceptive signals from the body, and helps generating adapted behavioral responses and learning about external threats.

## Acknowledgments

We thank Dr. Ayah Khubieh for help with the characterization of VGluT2^Cre^ mice, Heather Murray for support with mouse genotyping and histology, Dr. Michael Kintscher for help with the analysis of mouse behavior, Dr. Carmen Sandi for sharing equipment, Dr. Bernard Schneider and the EPFL Bertarelli Foundation Gene Therapy Platform for AAV vector packaging, and Dr. Nicolas Chiaruttini and Dr. Olivier Burri for their input with the ABBA-QuPath analysis pipeline. Images were acquired at the bio-optical imaging platform of EPFL (BIOP). The study was supported by grants from the Swiss National Science Foundation (SNF), 31003A_176332 / 1 and 310030_204587 / 1, to R.S.

## Figure Legends

## Materials and Methods

### Experimental model and subject details

All experimental procedures were performed with wildtype or genetically modified mice (*Mus musculus*) and were authorized by the veterinary office of the Canton of Vaud, Switzerland (authorizations VD3274 and VD3518). To limit variability in defensive behavioral responses arising from sex differences, noted in previous studies on defensive behaviors (Maren et al., 1994; Pryce et al., 1999; Gruene et al., 2015), only male mice were used for the experiments. The following mouse lines were used: 1) transgenic *CaMKII^Cre^* (B6.Cg-Tg(Camk2a-cre)T29-1Stl/J; JAX stock #005359; RRID:IMSR_JAX:005359; see Tsien et al., 1996); 2) transgenic *vGluT2^Cre^* (Slc17a6^tm2(cre)Lowl^/J; JAX stock #016963; RRID:IMSR_JAX:016963; Vong et al., 2011); 3) transgenic *Scnn1a^Cre^* mouse line (B6;C3-Tg(Scnn1a-cre)3Aibs/J; JAX stock #009613; RRID:IMSR_JAX:009613; see Madisen et al., 2010); 4) transgenic *Etv1^Cre^* mouse line, (B6(Cg)-Etv1^tm1.1(cre/ERT2)Zjh^/J; JAX stock #013048; RRID:IMSR_JAX:013048; Taniguchi et al., 2011); 5) Cre-dependent tdTomato reporter mouse *Rosa26LSL-tdTomato* (B6.Cg-Gt(ROSA)26Sor^tm9(CAG-tdTomato)Hze^/J; JAX stock #007909; RRID:IMSR_JAX:007909; also called “Ai9”; Madisen et al., 2010); 6) wild-type (WT) mice C57Bl/6J (JAX stock #000664; RRID:IMSR_JAX:000664). Transgenic mouse lines were back-crossed to a C57Bl/6J background for at least five generations. After weaning, male mice were group-housed under a 12/12 h light/dark cycle (lights on at 7am) with food and water *ad libitum,* and were separated into single cages starting one day prior to surgery until the end of experiment. For behavioral experiments (Figures 1, 5 and 6), mice from 1-2 litters were randomly assigned to control (GFP-expressing or saline controls) or effect group (Arch-or Halo-expressing).

## Method details

### Plasmid DNA cloning

We constructed a custom vector AAV:hSyn:FLEX:Synaptophysin-mCherry-IRES-eGFP:CW3SL, to drive the Cre-dependent co-expression of Synaptophysin-mCherry as a presynaptic marker, and a soluble cytosolic eGFP (construct “A”). To accommodate the IRES-based vector size to the packaging capacity of AAV, we chose a monomeric mCherry fluorophore as a tag for Synaptophysin, and used a reduced-size 3’UTR sequence known as W3SL (∼0.4 kb), consisting of a minimal WPRE3 sequence and SV40 late polyadenylation signal (Choi et al., 2014). Construction of the vector for AAV2/8 virus production was done using conventional cloning approaches based on PCR amplification and ligation of DNA fragments. In brief, a W3SL sequence was PCR amplified from the vector pAAV-CW3SL-EGFP (a gift from Bong-Kiun Kaang; Addgene #61463; RRID:Addgene_61463; see Choi et al., 2014). To investigate the effect of order of the expression cassettes relative to the IRES sequence, we also generated a swapped plasmid “B” (eGFP-IRES-Synaptophysin-mCherry sequence). We tested the relative expression of eGFP and Synaptophysin-mCherry in primary neuronal cultures transfected with the plasmid A or B using Ca^2+^ -phosphate method, and noted that the cassette downstream of the IRES sequence had a lower expression level (Osti et al., 2006). Therefore, for AAV2/8 packaging, we used the vector “A” which showed a higher synaptophysin-mCherry expression as compared to “B”. Intermediate and final DNA plasmids were verified by sequencing prior to the next cloning step or AAV packaging. The annotated plasmid sequence can be downloaded from the open data repository associated with the manuscript.

### Viral vectors and stereotaxic coordinates

For the optogenetic inhibition experiments (Figures 1), we expressed the inhibitory opsin archaerhodopsin-3 (Arch; Chow et al., 2010), by injecting AAV1:CAG:FLEX:Arch-eGFP (200nl; 2.05·10^12^ vg[vector genomes]/ml; University of North Carolina Chapel Hill vector core [UNC-VC], NC, USA) bilaterally into the pInsCx of *CaMKII^Cre^* mice. The stereotaxic injection coordinates were: ML (medio-lateral) ±4.1 mm, AP (anterior-posterior) -0.9 mm, DV (dorso-ventral) -3.85 mm; from the bregma point on the skull surface. Mice in the control group were injected with either an AAV1:CAG:FLEX:eGFP (200nl; 4.4·10^12^ vg/ml; UNC vector core), or an AAV1:shortCAG:FLEX:eGFP vector (200nl; 4.4·10^12^ vg/ml; University of Zürich viral vector facility [UZH-VVF], Switzerland; cat. v158-1).

To target Arch expression to LA-projectors of the pInsCx (Figure 5), we injected WT mice with a retrograde vector AAVretro:EF1α:mCherry:IRES:Cre (200nl; 1.30·10^13^ vg/ml; Addgene #55632-AAVrg) bilaterally into the anterior LA using the stereotaxic coordinates: ML ± 3.42 mm; AP -1.12 mm, DV -4.45 mm. Expression of Arch (or eGFP in control mice) in the LA-projecting neurons of the pInsCx was achieved by bilateral injection of the same Cre-dependent viral vectors in the pInsCx, as described above.

For optogenetically-assisted circuit mapping (Figures 3 and 4), we injected AAV8:EF1α:FLEX:Chronos-eGFP (200nl; 4.50·10^12^ vg/ml; UNC vector core) or AAV8:hSyn:Chronos-eGFP (200nl; 6.50·10^12^ vg/ml; UNC vector core) vectors bilaterally into the pInsCx (coordinates as above) of *CaMKII^Cre^* (Figure 4) or *CaMKII^Cre^* × *tdTomato* mice (Figure 3). When *CaMKII^Cre^* mice were used, an AAV8:CAG:FLEX:tdTomato (200nl; 6.5·10^12^ vg/ml; UNC vector core) was additionally injected into the LA (coordinates as above) in order to identify CaMKII^Cre^-positive pyramidal neurons by tdTomato fluorescence for subsequent patch-clamp recordings, as in the experiments using CaMKII^Cre^ × tdTomato double transgenic mice.

For the *in-vivo* Ca^2+^ imaging experiments (Figure 6), *CaMKII^Cre^* mice were injected with AAV8:hSyn:FLEX:GCaMP6m (200nl; 6.4·10^12^ vg/ml; University of Zürich VVF; cat. v290-1), into the left LA (coordinates as above). Additionally, these mice were bilaterally injected in the pInsCx (coordinates as above) with either an AAV8:hSyn:eNpHR3.0-eYFP vector (200nl; 7.3·10^12^ vg/ml; University of Zürich VVF; cat. v560-8) in the Halorhodopsin group, or with the same volume of phosphate buffered saline (PBS) in the control group.

For anterograde labeling of axonal projections (Figure 2, A-H), a custom-cloned vector AAV8:hSyn:FLEX:Synaptophysin-mCherry:IRES:eGFP (see above; packaged by Bertarelli Foundation Gene Therapy Platform at EPFL), was injected into the right pInsCx of *CaMKII^Cre^* mice (see coordinates above). Mice were processed for histological analysis of projection areas three weeks later.

For rabies-mediated retrograde transsynaptic tracing experiments (Figure 2, I-N), vGluT2^Cre^ mice were first injected into the right LA (stereotaxic coordinates: ML 3.42 mm; AP -1.12 mm; DV -4.45 mm) with a tricistronic vector AAV1:hSyn:FLEX:TVA:2A:eGFP:2a:oG (250 nl; 5.3·10^12^ vg/ml; cat. v243-1; University of Zürich VVF). Three weeks later, the rabies vector EnvA:ΔG:RV:dsRed (250 nl) was injected at the same coordinates, and the mice were processed for histological analysis seven days later.

### Stereotaxic surgery procedures

Stereotaxic injection of viral vectors and implantation of optic fibers (Figures 1 and 5) or GRIN lenses (Figure 6) were performed in 6-7 week-old male mice in a single surgery session, as previously described (Tang et al., 2020). Briefly, mice were anaesthetized with a gas mix of isoflurane in O_2_ (3% for induction and 1.5% for maintenance), and head-fixed to a stereotaxic apparatus (Model 942, David Kopf Instruments, Tujunga, CA, USA) using non-rupture ear-bars (Zygoma ear cups, Model 921, David Kopf Instruments). A mix of lidocaine and bupivacaine (∼50μl of 1 mg/ml and 1.25 mg/ml, respectively) in 0.9% NaCl saline was injected subcutaneously for local analgesia. The skull was exposed, and small craniotomies were drilled at the specific stereotaxic coordinates for virus injections and implantations. In case of implantations, an additional hole was drilled to insert an anchoring micro-screw (Cat#AMS90/1B-100; Antrin Miniature Specialties, Fallbrook, CA, USA). The viral suspension was injected using pulled glass pipettes and an oil hydraulic micromanipulator (MO-10, Narishige Group, Tokyo, Japan) at a speed of 100 nl/min; 5 min waiting time was allowed after each injection to prevent backflow.

Optic fiber implants for *in-vivo* optogenetic experiments (Figures 1 and 5) were custom-made from a 200 μm core / 0.39 NA / 230 μm outer diameter optic fiber (FT200EMT; Thorlabs Inc, Newton, NJ, USA) glued inside 1.25 mm diameter ceramic ferrules (CFLC230; Thorlabs) as described in (Sparta et al., 2012). During surgery, the implantable part (230 μm diameter optic fiber) was slowly advanced until the tip reached a position 500 μm above the injection site. For *in-vivo* electrophysiology recordings (Figure S1), an optrode was implanted above the injection site in a similar procedure; its ground wire was pre-soldered to an anchoring micro-screw.

For *in-vivo* Ca^2+^ imaging (Figure 6), a GRIN lens (500 μm / 6.1 mm ProView™ integrated lenses; cat. 1050-004413; Inscopix Inc., Palo Alto, CA, USA) was implanted with its tip 100 μm above the virus injection site in the anterior LA. To reduce tissue deformation, a 25G medical injection needle was first slowly inserted and retracted to mark the track for GRIN lens implantation. The GRIN lens was lowered with alternating down (200 μm) and up (50 μm) movements to minimize brain tissue deformation until 200μm from the intended DV coordinate.

After insertion of implants into the brain, their outer surfaces (ceramic ferrule, GRIN lens wall, the microscope docking platform, anchoring screw) as well as the skull bones were treated with a light-curing adhesive iBond Total Etch (Kulzer GmbH, Hanau, Germany) and secured in place by light curing dental cement (Tetric EvoFlow, Ivoclar Vivadent, Schaan, Liechtenstein). The open end of the GRIN lens at the integrated docking platform was sealed using Kwik-Sil (World Precision Instruments, Sarasota, FL, USA). The skin was stitched and covered with Betadine solution; the drinking water was supplemented with 1 mg/ml paracetamol, and the animals were monitored for the following 6 days to ensure adequate post-surgical recovery.

### Behavior testing

Behavioral experiments (Figures 1, 4-6) were conducted four weeks post-surgery using an auditory cued fear conditioning task. Mice were first habituated to the experimenter and to head tethering with the fiber-optic patch cord (for optogenetic experiments) or a dummy miniature - microscope (for Ca^2+^ imaging experiments) for 15 mins each on 5-6 consecutive days. Fear conditioning was conducted on 3 consecutive days in a conditioning chamber of a Video Fear Conditioning Optogenetics Package for Mouse (MED-VFC-OPTO-M, Med Associates Inc., Fairfax, VT, USA) under control of VideoFreeze® software (Med Associates Inc.). The conditioning chamber was located inside a sound-attenuated box (63.5 cm wide, 35.5 cm high, 76 cm deep; NIR-022MD, Med Associates Inc.) equipped with a speaker, and a CMOS video camera with a near-infrared filter for continuous video recordings of mouse behavior at 30 fps.

On day 1 (habituation), optic fiber patch cords, or an nVoke miniature - microscope (Inscopix Inc.) were attached to the ceramic ferrules or the GRIN lens docking platform, respectively, before placing one mouse at a time in the fear conditioning chamber (context A: rectangular plexiglass walls, metal grid floor, all pre-cleaned with 70% ethanol). Six tone blocks (CS), each consisting of 30 s long 1 Hz trains of tone beeps (100 ms long beeps of 7 kHz, 80 dB, 2 ms rise time) were presented at pseudo-random intervals. On day 2 (training), mice were placed in the same chamber and presented with six pseudo-randomly spaced CS blocks, each followed by an electric footshock (US; 0.6 mA AC, 1 s long) delivered by a shock generator (ENV-414S, Med Associates Inc.). On day 3 (retrieval), one mouse at a time was placed in a different chamber (context B: semi-circular walls, smooth acrylic floor, pre-cleaned with soap) and was re-exposed to four CS blocks to test fear recall.

During optogenetic inhibition experiments with Arch (Figures 1 and 5), green light (561 nm) was delivered on the fear conditioning day (day 2) for 3 s, starting 1 s before the footshock using a diode-pumped solid state laser (MGL-FN-561-AOM-100mW; CNI Lasers) via 200 μm core / 0.22 NA optic fiber patch cords (Doric Lenses Inc., Quebec, Canada). To avoid residual light leakage through the AOM module of the green laser, an additional mechanical shutter (SHB05T; Thorlabs) was introduced at the output before the fiber. The shutter was closed at all times except when laser light was applied. The laser power was adjusted for each animal to each a total light output power of 10 mW at the fiber tip.

### In-vivo electrophysiology

Custom-built optrodes were fabricated using an optic fiber (200 μm core / 230 μm outer diameter, NA 0.39; FT200EMT; Thorlabs) as described (Anikeeva et al., 2012; Tang et al., 2020). In brief, four tetrodes were twisted of insulated Pt/Ir wires (25 µm diameter; California Fine Wire, Grover Beach, CA, USA) and glued onto the external surface of the optical fiber inserted in the ceramic ferrule (CFLC230; Thorlabs), held in turn by the movable nut of a micro-drive (Axona Ltd, St Albans, UK). The free ends of the wires were attached to the pins of an NPD-18-VV-GS nano-connector (Omnetics Connector Corp., Minneapolis, MN, USA) that was fixed to the nut of the micro-drive. The impedance of recording channels was reduced to values typically <100 kΩ at 1 kHz in PBS by plating the Pt/Ir wire tips with platinum black solution (NeuraLynx, Bozeman, MT, USA) according to manufacturer’s manual.

One day before the start of the fear conditioning protocol, the optrode was advanced by ∼100-300 μm in the ventral direction using a micro-drive until a target depth of 3.7 mm DV was reached, while the mouse was under ketamine anaesthesia.

The AP activity of pInsCx neurons in freely moving mice was recorded at 40 kHz sampling frequency with a 16-channel amplifier ME16-FAI-mPA under control of the MC_Rack software (Multi Channel Systems, Reutlingen, Germany). On day 2 of the fear conditioning protocol, 3 s long laser light pulses (561nm, 10 mW) were applied starting 1 s before every second US. The spiking activity of units in the pInsCx was compared to that during footshocks without light pulses, to estimate the efficiency of Arch in silencing US responses in the pInsCx. TTL pulses triggering the laser (from Master-8 pulse generator; A.M.P.I, Jerusalem, Israel) and those marking CS and US stimuli (from VideoFreeze), were co-sampled by the recording amplifier.

### Miniature - microscope Ca^2+^-imaging

The cellular Ca^2+^ dynamics during a 3-day auditory cued fear conditioning was imaged in the LA of freely moving mice (Figure 6) using an nVoke system (Inscopix Inc.; see Stamatakis et al., 2018). Throughout the behavioral sessions, the GCaMP6m fluorescence was continuously imaged at a rate of 30 fps using blue LED light (450 nm) to excite GCaMP6m. The electronic focusing system of the miniature - microscope cycled through three focal planes spaced by 75-100 μm, leading to an effective sampling frequency of 10 fps per plane. Synchronization between the behavioral -and the Ca^2+^ -imaging video streams was achieved by recording CS and US presentations with the Inscopix GPIO TTL inputs.

For silencing of incoming pInsCx axons in the LA (Figure 6 and Figure S6), Halorhodopsin (eNpHR3.0) expressed by pInsCx axons was activated by a red-shifted LED of the nVoke microscope (630 nm), switched on 1s before the start of every second US (footshocks #1, 3 and 5), for a period of 3s (see Figure 6B). The Ca^2+^ signals during these US presentations were compared to those during the footshocks #2, 4 and 6 to estimate to the effect of optogenetic inhibition.

### Slice electrophysiology

*Ex-vivo* electrophysiology in slices (Figures 3 and 4) was performed 4 - 5 weeks following the injection of AAV vectors driving the expression of Chronos in the pInsCx (see above). One mouse at a time was deeply anaesthetized with isoflurane (3% in O_2_), decapitated, and 300 μm thick coronal brain slices containing the LA were made with a Leica VT1200S vibrating blade microtome (Leica Microsystems, Wetzlar, Germany). Slicing was done in ice-cold N-methyl-D-glutamine (NMDG) based slicing buffer (Ting et al., 2014), containing (in mM): 110 NMDG, 2.5 KCl, 1.2 NaH_2_PO_4_, 20 HEPES, 25 Glucose, 5 Na-ascorbate, 2 thiourea, 3 Sodium pyruvate, 10 MgCl_2_, 0.5 CaCl_2_, saturated with carbogen gas (95% O_2_/5% CO_2_), pH 7.4, adjusted with HCl. All chemicals were from Sigma-Aldrich (Merck KGaA, Darmstadt, Germany), except HEPES, NaCl, KCl (which were from Thermo Fisher Scientific, Waltham, MA, USA), and MgCl_2_ (AppliChem, Darmstadt, Germany). Slices were stored for 7 min at 36°C in a chamber containing slicing buffer, and then transferred into a chamber containing carbogen-saturated storage solution (at room temperature), containing (in mM): 92 NaCl, 2.5 KCl, 30 NaHCO_3_, 1.2 NaH_2_PO_4_, 20 HEPES, 25 glucose, 5 Na-ascorbate, 2 Thiourea, 3 Na-pyruvate, 2 MgCl_2_ and 2 CaCl_2_, pH 7.4 (Ting et al., 2014). Acute slices containing the amygdala (300 µm thickness) were sorted according to anterior - posterior position; we mainly used the anterior - most slice, and the second anterior - most slice for recordings (see Figure S3D). Whole-cell patch-clamp recordings were performed with a standard extracellular solution containing (in mM): 125 NaCl, 2.5 KCl, 25 NaHCO_3_, 1.2 NaH_2_PO_4_, 25 glucose, 0.4 Na-ascorbate, 3 Myo-Inositol, 2 Na-pyruvate, 1 MgCl_2_ and 2 CaCl_2_, pH7.4, saturated with carbogen gas. The patch-clamp recording set-up was equipped with an upright microscope (BX51WI; Olympus, Tokyo, Japan) with a 60x / 0.9 NA water-immersion objective (LUMPlanFl, Olympus) and an EPC10 USB double patch-clamp amplifier (HEKA Elektronik GmbH, Reutlingen, Germany).

For the activation of Chronos and Arch in slice experiments (Figures 3,4; Figure S1A-G; Figure S3), high-power LEDs (CREE XP-E2, 460 and 530 nm respectively; Cree Inc., Durham, NC, USA) were custom-coupled into the epifluorescence port of the microscope using a 30 mm cage optic mounting system and focusing lenses (Thorlabs). The 530 nm LED was also used for excitation of tdTomato in brain slices. Illumination power and timing was controlled by the EPC10 amplifier connected to the LED driver (BLS-1000-2, Mightex Systems, Toronto, Canada). Irradiance was measured with a fast photodiode (DET36A/M, Thorlabs) in the illumination light path, whose readings were digitized and calibrated using an optical power meter (1918-R with 818-UV detector; NewPort, Irvine, CA, USA) under the 60x objective.

Patch-clamp recordings for optogenetically-assisted circuit mapping experiments in Figure 3, A-F and Figure S3 were done at room temperature (22-24°C), while all the other patch-clamp recordings were done at near-physiological temperature (34°C). In the latter case, the temperature was controlled by an in-line heater SHM-6, a heated recording chamber RC-26GL/PM-1 and a thermostatic control unit TC-344B (all from Warner Instruments, Holliston, MA, USA).

For the whole-cell patch clamp recordings in Figure 3A-F, and in Figure 4, a Cs^+^-based pipette solution was used, containing (in mM): 140 Cs-gluconate, 10 HEPES, 8 TEA-Cl, 5 Na-phosphocreatine, 4 Mg-ATP, 0.3 Na-GTP, 5 EGTA, pH 7.2 adjusted with CsOH. oEPSCs were evoked by 1 ms blue light pulses (∼ 5 mW/mm^2^) in the presence of the GABA_A_ receptor antagonist gabazine (SR-95531, 5 μM; Abcam, Cambridge, UK), at 34°C. The oEPSC amplitudes were quantified from the average of 10 sweeps. The AMPA-EPSC was measured as the peak of the inward current at -70 mV holding potential, and the NMDA-EPSC was measured as the outward current amplitude between 48-50 ms after the onset of the light pulse.

The remaining whole-cell patch clamp recordings (Figure 3G-I; Figure S1A-G) were made using a K^+^-based pipette solution containing (in mM): 145 K-gluconate, 8 KCl, 10 HEPES, 3 Na-phosphocreatine, 4 Mg-ATP, 0.3 Na-GTP, 5 EGTA, pH 7.2 adjusted with KOH.

### Immunohistochemistry and histological analysis

For the *post-hoc* histological analysis of opsin expression and implant placements following behavior (Figure S1L, Figure S5), and for the anatomical experiments (Figure 2), the mice were euthanized by a lethal injection of pentobarbital (150 mg/kg body weight), then transcardially perfused with 4% paraformaldehyde (PFA; Sigma-Aldrich) solution in PBS. The brains were post-fixed overnight in 4% PFA and dehydrated in 30% sucrose solution in PBS. Serial coronal sections (40 μm) were cut from frozen brains with a HM450 sliding microtome (Thermo Fisher Scientific, Waltham, MA, USA).

For the mapping of output projections using the Synaptophysin-mCherry construct (Figure 2A-H), immunohistochemistry was performed with a rabbit anti-RFP antibody (ab62341; Abcam; RRID:AB_945213; 1:400 dilution), and a secondary, donkey anti-rabbit Alexa-568 antibody was used (A10042; Thermo Fisher Scientific; RRID:AB_2534017; 1:200 dilution). For immunohistochemical staining of CaMKII+ neurons in the LA (Figure S2 B-D), immunohistochemistry was performed with a mouse monoclonal anti-CaMKII primary antibody (Cat# 10011437, Cayman Chemical, RRID:AB_10352211; 1:500 dilution), followed by a goat anti-mouse Alexa 555 antibody (A-21424, Thermo Fisher Scientific, RRID:AB_141780; dilution 1:1000).

Free-floating brain sections were mounted on Superfrost® Plus slides (Thermo Fisher Scientific) and embedded in Fluoroshield® mounting medium containing DAPI (Sigma-Aldrich). The sections were imaged with a slide scanner automated microscope VS120-L100 (Olympus) with a 10x/0.4 NA objective, or with an upright Leica SP8 confocal microscope (Leica Microsystems).

## Quantification and statistical analysis

### Behavior data analysis

Video recordings of the animal behavior (30 fps) were analyzed using ezTrack software (Pennington et al., 2019) to generate a movement index trace. The space above the mouse containing optic patch cords or the cable of the miniature microscope were cropped to minimize movement artifacts. The movement index trace was then used to compute a binary freezing trace using custom procedures in IgorPro 7 (WaveMetrics Inc., Lake Oswego, OR, USA). The animal was considered immobile (“freezing”) if the movement index for at least 0.5 s was below a threshold of 40 or 120 arbitrary units (a.u.) without or with cable attachment, respectively. The trace of percent time spent freezing (binned at 10 s) was calculated as a time-average of individual binary freezing state traces averaged across mice in each group. The CS-specific freezing was quantified as the percentage freezing during the 30s CS-presentation *minus* the freezing during the 30s no-CS period (shown as light grey bars, see e.g. Figure 1G). The experimenter and data analyst were blinded to the assignment of each mouse to the control or test group.

### Data analysis for *in-vivo* unit activity

For the analysis of 16-channel *in-vivo* extracellular recording data, the raw data was converted from MC_Rack into HDF5 format using Multi Channel Data Manager software (MultiChannel Systems). The data was then processed using custom-written routines in IGOR Pro 7 (WaveMetrics, USA), except the spike clustering step. Briefly, for the spike detection, the voltage traces were band-pass filtered (0.6-6 kHz; 4th-order Butterworth filter). Footshock stimulation artefacts were blanked by zeroing under manual control. Negative amplitude spikes were detected by thresholding (typically set at −3.8 standard deviations), and the spike location was determined by the largest event within each tetrode. Individual spike cut-outs (filtered at 0.3-6 kHz with a band-pass 4th order Butterworth filter) were subjected to clustering analysis using the MClust toolbox (Dr. David Redish; University of Minnesota, USA) in MATLAB (MathWorks, USA), which applied an unsupervised clustering algorithm KlustaKwik (Rossant et al., 2016) using the spike valley and the principal components PC1-PC3 as parameters. The clustering output was manually quality-controlled by checking the average spike waveform similarity and visualizing the cluster projections; occasionally, several clusters were fused together if they were not well separated. Next, the isolation distance (ID) and L-ratio were computed to control for type I and type II errors (Schmitzer-Torbert et al., 2005). Only clusters with ID > 24 and L-ratio < 0.5 were retained for further analysis; this reduced the total number of clusters in the sample from n = 54 to 30 in N = 2 mice. The clustered data were re-imported into IgorPro for subsequent waveform matching, frequency and Z-score calculation, alignments to the stimuli and plotting. The units recorded on day 2 were classified as US-responsive if the average Z-score exceeded a value of 2 during the footshock, calculated from the control (no light) US presentations #2, 4 and 6.

### Data analysis for slice electrophysiology

Data was exported from the PatchMaster software (HEKA Elektronik), and analyzed in IgorPro 7 (WaveMetrics) using custom scripts.

### Ca^2+^-imaging data analysis

The initial processing steps for data analysis were done using the built-in analysis pipeline of the Inscopix Data Processing Software (IDPS; Inscopix Inc.). This included: 1) deinterleaving of the video streams into the individual focal planes; 2) spatial filtering; 3) motion correction; 4) export of the synchronization timestamps. Motion-corrected videos (one per each of three focal plane) were exported as multiplane TIFFs and processed using an automated CaImAn pipeline (Giovannucci et al., 2019) for source separation and ROI assignment based on constrained non-negative matrix factorization (Pnevmatikakis et al., 2016; Zhou et al., 2018). The output from CaImAn analysis was imported into IgorPro 7 for further processing using custom-written routines. The timestamps for every data point, as well as the synchronization signals for alignment with the camera monitoring the behavior, were imported from IDPS. Duplicate cells appearing on neighboring focal planes were removed semi-automatically based on the lateral proximity of their centroids and on the cross-correlation of fluorescence intensity traces. The intensity traces for each i^th^ cell (𝐹𝐹_𝑖𝑖_(𝑡𝑡)), were standardized by converting them to Z-score traces as 𝑍𝑍_𝑖𝑖_(𝑡𝑡)=(𝐹𝐹_𝑖𝑖_(𝑡𝑡)−〈𝐹𝐹_𝑖𝑖_(𝑡𝑡)〉)/𝜎𝜎(𝐹𝐹_𝑖𝑖_(𝑡𝑡)), where 〈𝐹𝐹_𝑖𝑖_(𝑡𝑡)〉 and 𝜎𝜎(𝐹𝐹_𝑖𝑖_(𝑡𝑡)) are, respectively, the mean and the standard deviation of the i^th^ cell intensity, calculated from the whole duration of the experiment.

The onsets of US presentations were used to align the excerpts of 𝑍𝑍_𝑖𝑖_(𝑡𝑡) traces, which are represented by individual rows on the color-coded raster plots (see Figure 6E as an example). Cells were classified as US-responders, if they had an average Z-score > 1.0 during 2.5 s period following the onset of US presentations #2, 4, and 6. The average Z-score values were used to sort the color-coded raster plots in the descending order of the response strength (Figure 6E and Figure S6C). The difference between the Ca^2+^ transient amplitude in the absence *minus* the presence of light (Δ z-score^US^ = z-score^US, light^ -z-score^US, no light^) was calculated for all US-responders (Figure 6H). Their distribution was fitted to a double Gaussian function, with the same initial parameters for both control and halorhodopsin groups, and the position of the first peak and the y-offset were constrained to zero.

### Histological image analysis

Histological image analysis was performed on every second coronal section; i.e. at 80 μm distance, for the brain volume containing an implant (in case of *post-hoc* analyses), or in the range from the prefrontal cortex up to the end of the cerebellum (in case of anatomical experiments; Figure 2). For post-hoc analyses, the vectorized brain atlas maps (Franklin and Paxinos, 2016) were fitted to the image by affine transformation in Adobe Illustrator (Adobe, Mountain View, CA, USA). For the anatomical experiments (Figure 2), the slide scanner images were first imported into QuPath software (Bankhead et al., 2017) and then registered to the reference Allen Brain Atlas (2014) using a FIJI plugin ABBA, developed by the bioimaging platform at EPFL (Chiaruttini et al., 2022). Transformed atlas maps were next sent to QuPath, which performed automated detection of dsRed labelled cell bodies or synaptophysin-mCherry expressing synaptic terminals. Finally, the distribution of detected objects across atlas-defined regions throughout the whole brain was quantified using a custom Python script, averaged across animals and plotted using IgorPro scripts (https://github.com/bmi-lsym/ABBA-QuPath-post_processing), and the locations of detected cells were rendered into a 3D brain model (Figure 2L) using the Python-based Brainrender package (Claudi et al., 2020).

### Statistical analysis

Statistical testing was performed in GraphPad Prism 9 (GraphPad, San Diego, CA, USA). Before choosing the main statistical test, the distribution of the data was tested for normality using a Shapiro-Wilk test. If confirmed, a paired or unpaired version of the two-tailed Student’s t-tests was used for two-sample datasets. In case the data were not conformed to the normal distribution, two-tailed non-parametric tests were used: a Wilcoxon matched-pairs signed-rank test for paired comparisons, or Mann-Whitney U test for unpaired comparisons. When more than two comparison groups were present, we used either one-way ANOVA followed by Bonferroni post-hoc tests, or Kruskal-Wallis test followed by Dunn’s post-hoc test for multiple comparisons, in case of normally or non-normally distributed datasets, respectively. Post-hoc tests were only performed when the main test (ANOVA or Kruskal-Wallis) reported significant group differences (p < 0.05). For datasets influenced by two factors (e.g. group and time), we used a repeated-measures two-way ANOVA (RM-ANOVA) separately for each day of the fear conditioning protocol. If RM-ANOVA reported significance for the group factor or its interaction with the time factor, it was followed by Bonferroni post-hoc tests for multiple comparisons, to test the influence of group and/or time.

The statistical test used for each experiment is stated in the Results along with the statistical test summary (values of test statistics, degrees of freedom, p-values). Statistical significance is indicated in the Figures using asterisks according to the convention: p ≤ 0.05 (*), p ≤ 0.01 (**) and p ≤ 0.001 (***).

## Supplemental Information Titles and legends

**Figure S1.**
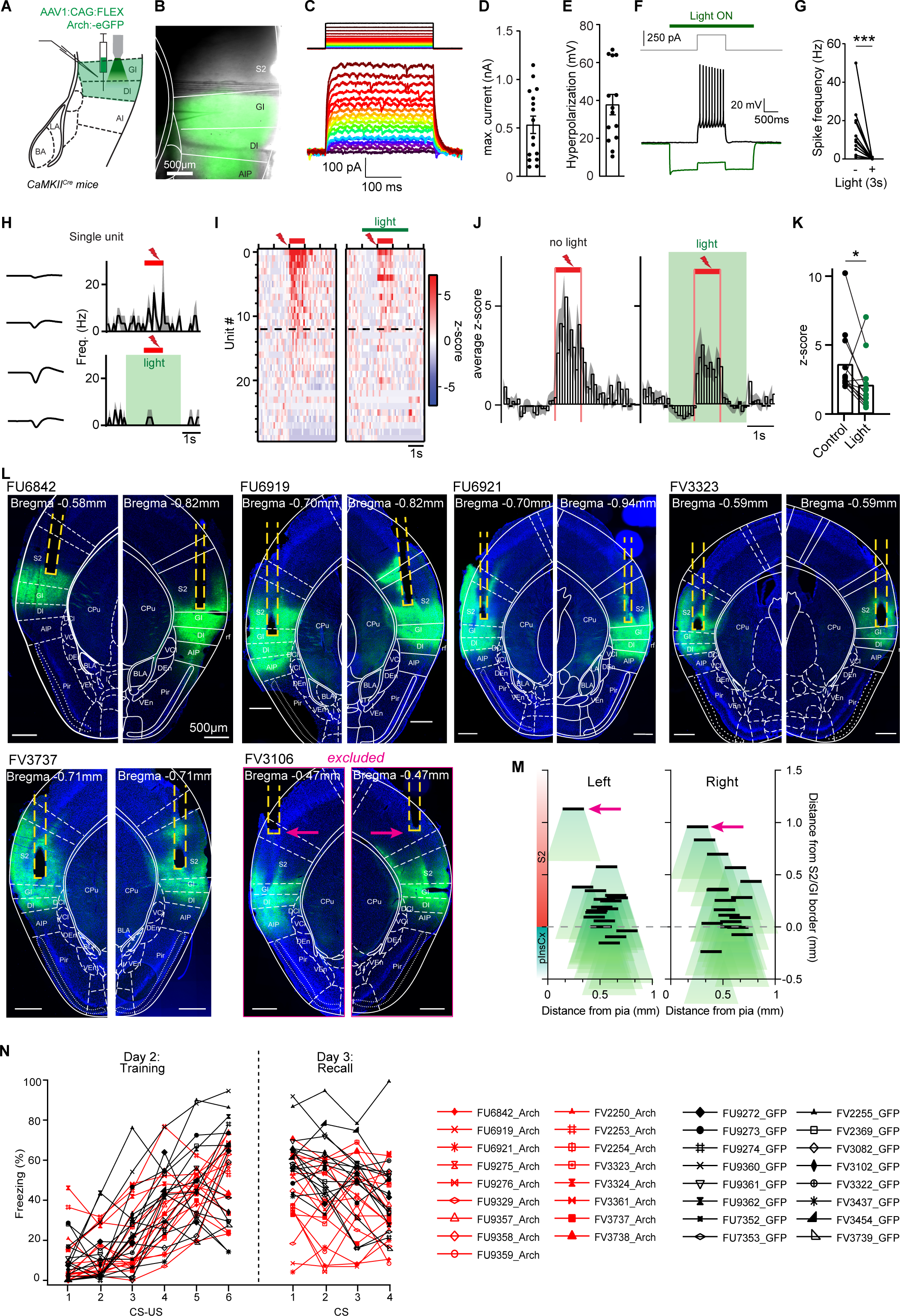
Control experiments for Arch - mediated optogenetic silencing, and post-hoc validation of Arch expression and fiber placements. Related to Figure 1. (A) Scheme of the experimental approach. (B) Expression of Arch-eGFP visualized in an acute slice of the pInsCx after excitation with 470 nm light. (C) Light-induced currents upon green light (530 nm) applications of increasing intensity (from 0 - 15 mW/mm^2^). (D) Maximal current amplitude measured in n = 16 recordings from N = 3 Arch - expressing mice. An average maximum current of ∼500 pA across neurons was observed. (E) Light-induced maximal hyperpolarization measured in n = 14 recordings in current clamp, starting at a membrane potential (V_m_) of ∼ -70 mV. (F) The AP-firing of an example neuron resulting from a current injection of +250pA was abolished by the application of a 3s-long green light pulse of maximal intensity (15 mW/mm^2^). (G) Across all recorded neurons (n = 12 in N = 3 mice), AP-firing was strongly suppressed by the application of green light (p < 0.005; W= -78, Wilcoxon test). *(H-K)* Arch was expressed uni-laterally in the pInsCx of CamKII^Cre^ mice (AAV1:CAG:FLEX:Arch-eGFP), and an optrode was implanted in the pInsCx. Three weeks later, the mice underwent the regular fear learning protocol, with n = 6 footshocks applied during the training session. Green light was applied during footshocks #2, 4, 6. (H) Example of a single recorded unit. *Left*, AP-waveforms recorded at each electrode of one tetrode. *Right*, AP-firing response (average ± SEM, n = 3 each) to footshocks without light (*top*), and with 3s green light application (*bottom*). (I) Color-coded plot of z-scored AP frequencies in response to n = 3 footshock applications without and with green light (*left* and *right*). The units were sorted according to the strength of the response in the absence of light (*left*); twelve units with a significant response were found (see dashed line). Note the decrease in response (except for one unit) when green light was applied to activate Arch. (J) Average z-scored AP-frequency for the n = 12 units in (I) without, and with light application (*left*, and *right*). (K) Individual data points, and averages (bars) of the peak z-scored AP-firing response to footshocks in the absence (*left*) and presence of green light (*right*). The peak AP-firing response was significantly reduced by green light (n = 12 footshock-responsive units recorded in N = 2 mice; p = 0.034, Wilcoxon’s test). (L) Post-hoc validations of Arch-eGFP expression for the experiments of Figure 1. The dataset contained N = 18 mice expressing Arch, and N = 16 mice expressing eGFP. For space reasons, we show a sub-set of the Arch-expressing mice. Each mouse is identified by a code (e.g. “FU6842” for the top-left mouse). For each mouse, we show the brain section in which the center of the fiber tip was located; fibers are enhanced by yellow dashed lines. Mouse brain atlas images (Franklin and Paxinos, 2016) were overlaid, and the Bregma value of the Franklin and Paxinos map is indicated for each side of the brain. Abbreviations for brain area names are as in (Franklin and Paxinos, 2016; and see Table 1). One mouse was excluded, because the fibers were clearly implanted outside the pInsCx (FV3106), leading to a final dataset of N = 18 Arch expressing mice. (M) Reconstruction of the placement of fiber tips (black lines) with respect to medio-lateral position (distance from pia, x-axis) and dorso-ventral position as estimated by distance from the S2 - pInsCx border (y-axis). A cone with a length of 500 µm was added below each fiber, to illustrate the assumed extent of silenced brain tissue. This assumes that light with an intensity of 10 mW at the fiber tip, can activate Arch over a distance of 500 µm, given that Arch is expressed in that area (Aravanis et al., 2007; Baleisyte et al., 2022). The position of the fiber tips in the excluded mouse (FV3106, see panel L) is indicated by purple arrows. (N) Freezing values (percent time spent freezing during 30s CS intervals) during the training session (*left*) and during the recall session (*right*), are shown for individual mice in the eGFP data set (black data, N = 16 mice), and for individual mice in the Arch data set (red data, N = 17 mice after exclusion of FV3106). These values are the individual data points for the average traces shown in Figure 1F, *left* (training session), and Figure 1H, *left* (recall session). The correspondence of each symbol to its mouse code is given (*right*).

**Figure S2.**
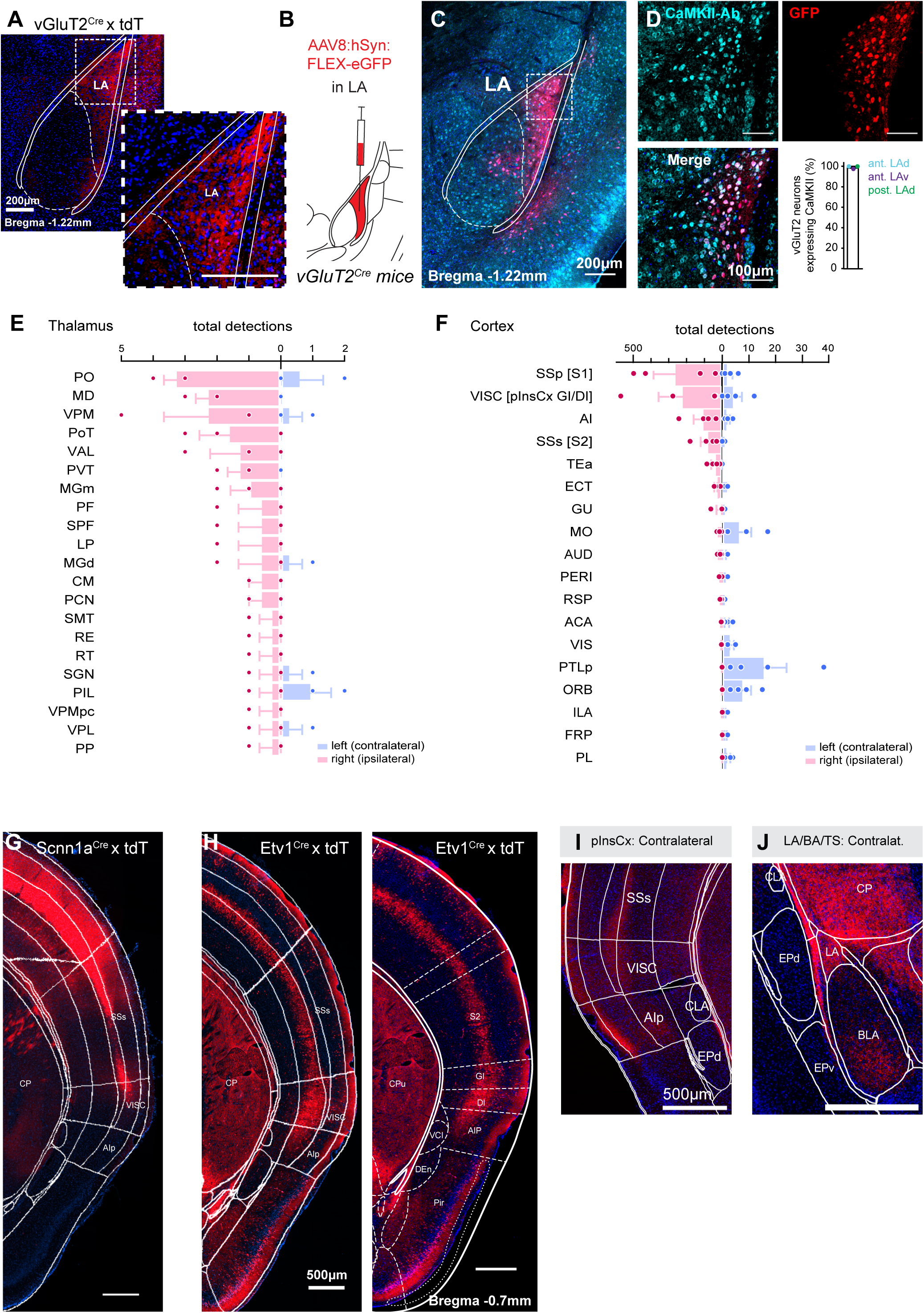
Suitability of a VGluT2^Cre^ mouse for rabies experiments in the LA; results from monosynaptically-restricted backlabeling; and additional data from further Cre mouse lines regarding the S2 - pInsCx border. Related to Figure 2. (A) Because the pInsCx makes a strong projection to the tail striatum adjacent to the LA (Figure 2D), it was necessary to identify a Cre mouse line that allows selective back-labeling from the LA. Based on the Allen Mouse Brain Atlas, we found that VGluT2 is expressed in the LA (data not shown; see https://mouse.brain-map.org/). To validate this, we crossed *VGluT2^Cre^* mice with a tdT reporter mouse, *Rosa26LSL-tdTomato*. This revealed tdTomato - positive cell bodies and axons in the LA (red channel), but no Cre-expression in the tail striatum adjacent to the LA. *Red*, tdTomato channel, *blue*, DAPI channel. (B-D) Cre-dependent expression of eGFP from an injected AAV8:hSyn:FLEX:eGFP expression vector (B) shows that Cre - expressing neurons are CamKII - positive principal neurons. eGFP fluorescence (red channel), and anti-CamKII-immunohistochemistry (cyan channel; C), and zoom-ins (D) in the area indicated by a box in (C) are shown. Note that essentially every Cre - expressing neuron is CamKII - positive (D, bottom panel, *left*). This is confirmed by quantification (D, bottom panel, *right*; n = 82/96 vGluT2/CaMKII neurons in the anterior - dorsal LA; n = 68/79 neurons in the anterior - ventral LA; and n = 91/101 neurons in a more posterior dorsal area of the LA; N = 1 mouse). (E, F) Average number of detected, back-labelled neurons in the thalamus (E), and in cortical areas comprising isocortex as defined in ABA (F). Data are mean ± SEM from N = 4 mice. The brain area abbreviations follow the ones of the ABA, except abbreviations given in brackets, which refer to the names used in this paper (see also Table 1 for a list of abbreviations). (G) *Scnn1a^Cre^* mice, which drive Cre - expression in layer 4 of sensory cortices (Madisen et al., 2010), were crossed with *Rosa26LSL-tdTomato* mice; coronal brain section images were then registered to the ABA with the ABBA tool (Materials and Methods). Note the localization of tdTomato - positive (thus, Scnn1a - positive) neurons on the border between the ventral S2, and the pInsCx - GI (dorsal part of “VISC”), thus defining the end of layer 4 in the pInsCx-GI, as expected. (H) *Etv1^Cre^* mice, which drive Cre - expression in layer 5 neurons (Yoneshima et al., 2006; Sürmeli et al., 2015), were crossed with *Rosa26LSL-tdTomato* mice. Note the labeling of upper layer 5 neurons in the S1 and S2, and the expansion of an Etv1 - positive neuron layer more ventrally, coinciding with the dorsal pInsCx-GI, and with the thinning of layer 4 in the pInsCx - DI (ventral part of “VISC”; *left* panel). In the *right* panel, the same image was aligned to the brain atlas of (Franklin and Paxinos, 2016). This illustrates that “VISC” in the ABA roughly corresponds to the granular, and dysgranular part of the pInsCx in Franklin and Paxinos (pInsCx GI and - DI, respectively). (I, J) Example images from the anterograde labeling starting in the pInsCx (see Figure 2A-H), for contralateral brain areas. Note the Synaptophysin-mCherry positive axons in superficial layers of S2, pInsCx GI and -DI contralaterally, and in the posterior agranular insular cortex (AIp; panel I), as well as in the contralateral LA, BA and tail striatum (J).

**Figure S3.**
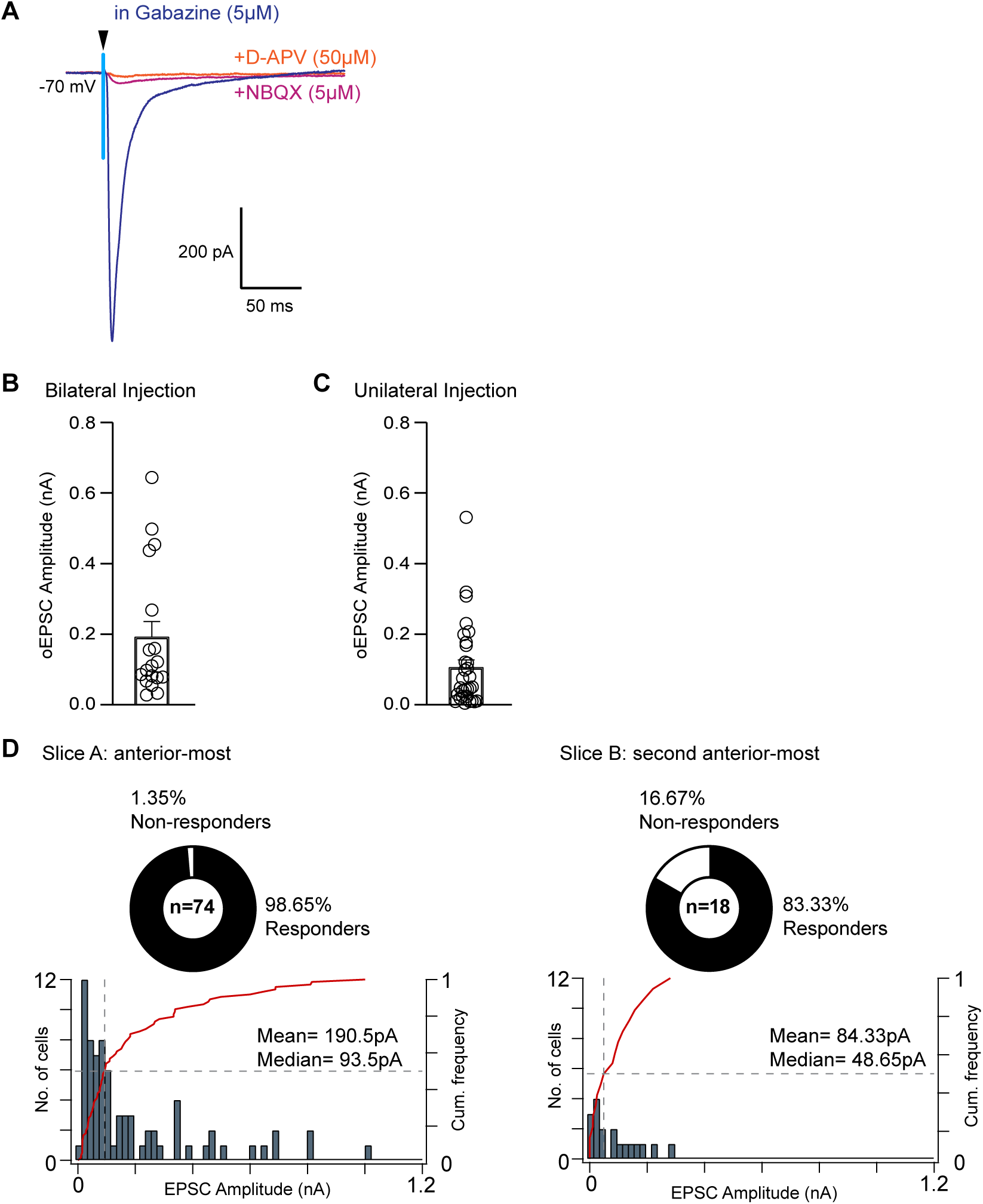
oEPSCs at the pInsCx - LA synapse are NBQX sensitive, and quantification of connection probability across the anterior - posterior axis in the LA. Related to Figure 3. (A) Example trace of an oEPSC recorded in a LA - principal neuron after expression of Chronos in the pInsCx, in the presence of Gabazine (5 µM) to block feedforward inhibition. Application of NBQX (5 µM) reduces the oEPSC to 5 % of its initial amplitude; application of AP5 (50 µM) further reduces the remaining oEPSC. (B) oEPSCs measured after bilateral expression of Chronos in the pInsCx, in a Cre-<u>in</u>dependent manner in *CamKII^Cre^* mice (using AAV8:hSyn:sChronos-eGFP vector). The resulting oEPSC amplitudes (191.8 ± 43.96 pA; n = 18 recordings from N = 4 mice), were indistinguishable from oEPSC amplitudes after bilateral Cre-dependent expression of Chronos in the pInsCx of *CamKII^Cre^* mice (Figure 3I; p = 0.146; U= 151, Mann Whitney U test). Thus, the Cre-dependent expression of Chronos in *CamKII^Cre^* mice did not lead to the exclusion of a sizable pool of CamKII - negative LA - projectors in the pInsCx. Therefore, we conclude that most LA - projectors are CamKII - positive neurons. (C) oEPSCs measured after expressing Chronos in a Cre-<u>in</u>dependent manner in *CamKII^Cre^* mice, by injecting the expression vector (AAV8:hSyn:Chronos-eGFP) unilaterally in one pInsCx (subsequent recordings were made in the ipsilateral LA). Note the smaller amplitude of oEPSCs (105.4 ± 21.92 pA, n = 30 recordings in N = 11 mice), as compared to the data set with bilateral injection (panel B; p = 0.0035; U= 171, Mann Whitney U test). This finding is consistent with the anatomical finding that pInsCx principal neurons project to both the ipsi- and contralateral LA (Figure 2, Figure S2I, J). (D) Analysis of connection probability, and of oEPSC amplitude depending on anterior - posterior position in the LA. Distributions of oEPSC amplitudes, and plots of the connection probability are shown for the anterior-most slice (*left*), and for the second anterior-most slice containing the LA (*right;* see Materials and Methods), for oEPSCs recorded after uni- and bilateral expression of Chronos in the pInsCx (see data sets in Figure 3I, and in panel B). The connection probability was higher in the anterior-most slice than in the second anterior-most slice (98.7% versus 83%; p = 0.022; Fischer’s exact test). Similarly, there was a trend towards a larger oEPSC amplitude in the anterior-most slice as compared to the second anterior-most slice, but this comparison did not reach statistical significance (p = 0.052, U= 468.5, Mann Whitney U test). These findings are consistent with the anatomical findings showing that pInsCx axons mainly innervate the anterior LA (Figure 2H).

**Figure S4.**
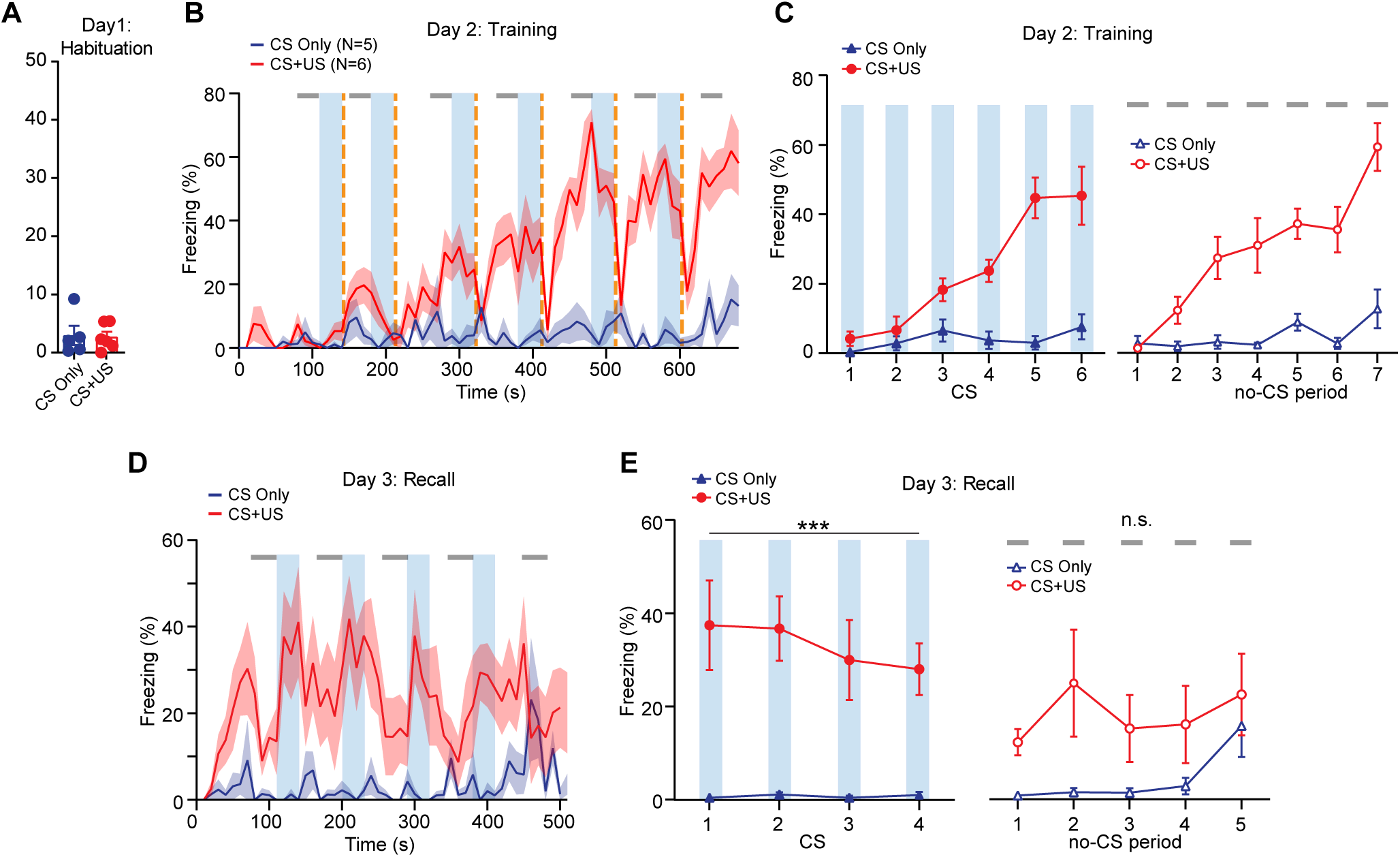
Behavior in the two groups of mice used for the measurement of AMPA/NMDA ratio changes after fear learning. Related to Figure 4. The behavioral data for the two groups of mice of Figure 4 (“CS-only”, and “CS-US”) was analyzed as in Figures 1, 5, where results from regular fear learning protocols are shown. (A) Average time spent freezing during the habituation day of each group of mice (CS-only, N = 5; CS+US, N = 6; blue and red data points). (B) The average time spent freezing during the training session for both groups, displayed at a time resolution of 10 s. In the CS+US group (which corresponds to the regular fear learning protocol; red trace; mean ± SEM for N = 6 mice), a gradual build-up of freezing was observed. Conversely, in the CS-only group in which no footshocks were applied, freezing remained low throughout the session, as expected (blue trace; mean ± SEM for N = 5 mice). (C) Analysis of time spent freezing (%) during the CS periods (*left*) and the 30-s no-CS periods (right; see gray bars in D), for both groups of mice during the training session. (D) Freezing behavior at 10s time resolution during the fear memory recall session, when 4 CS epochs (light blue stripes) were applied in a new context. Note the high levels of freezing in the CS+US group, that were additionally increased during the CS (red trace). Conversely, mice in the CS-only group showed low freezing. (E) Analysis of freezing during the 30-s CS times (*left*), and during 30-s no-CS periods immediately preceding each CS, and for one late no-CS period (see grey lines in panel D). The CS-induced freezing was significantly higher in the CS+US group as compared to the CS-only group (p=0.0001; F(1,9)=42.7, 2-way repeated measures ANOVA), but not during the no-CS times (p = 0.113, F(1,9)= 3.086; 2-way repeated measures ANOVA).

**Figure S5.**
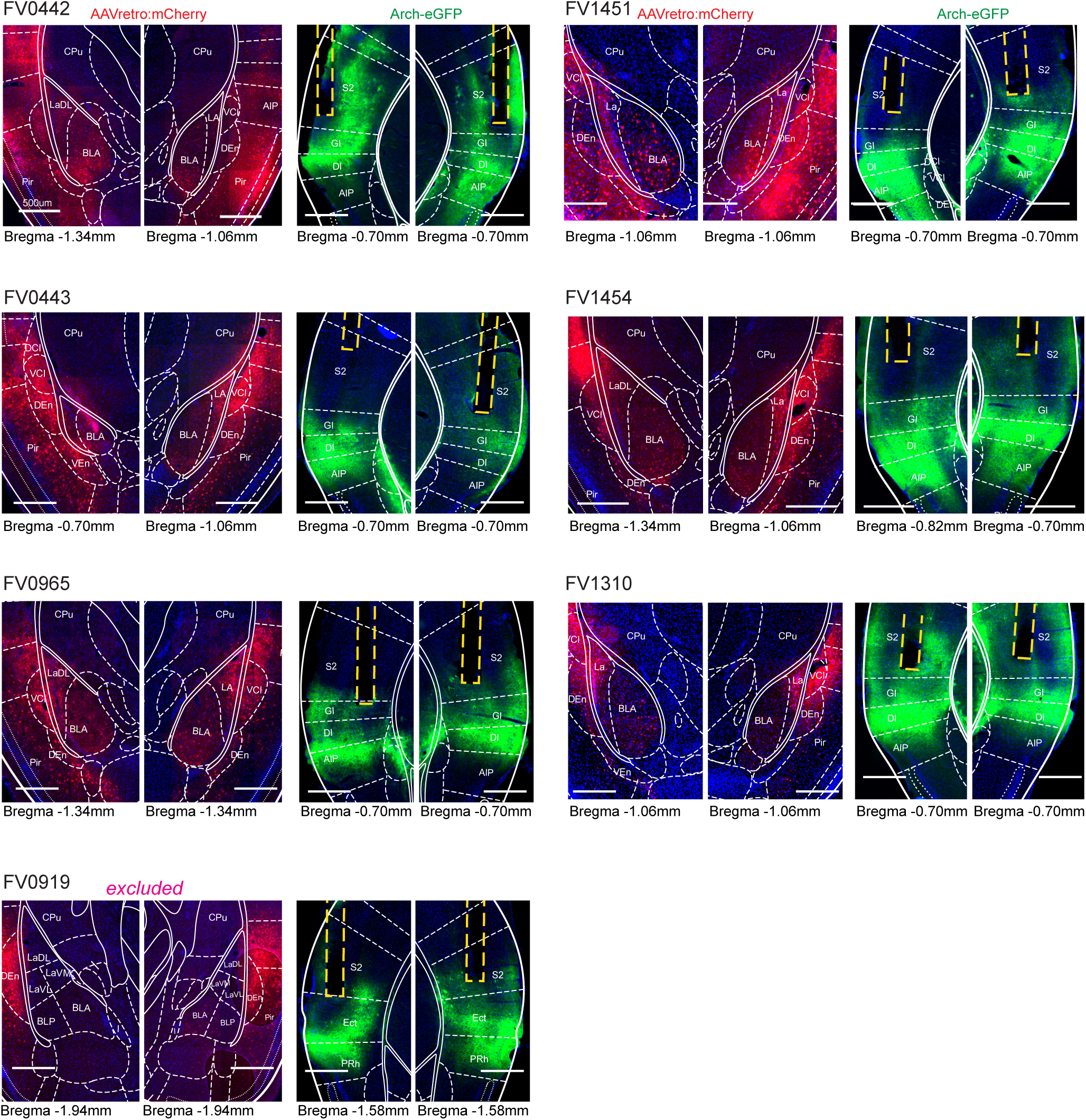
Post-hoc histological validations of AAVretro:mCherry and Arch-eGFP expression, and bilateral fiber placements in the pInsCx. Related to Figure 5. The images show the expression of mCherry driven by the injection of AAVretro: EF1α:mCherry:IRES:Cre into the left and right LA (panels with red fluorescence, *left*), and the expression of Arch-eGFP driven by the injection of AAV1:CAG:FLEX:Arch-eGFP into the left and right pInsCx (panels with green fluorescence, *right*); fiber positions are indicated by yellow dashed lines. Each mouse is identified by its code (e.g. FV0442, *top*-*left*). Maps from (Franklin and Paxinos, 2016) are overlaid; see Table 1 for the meaning of abbreviations for brain structure names. One mouse was excluded from the behavioral dataset, after we found that mCherry expression was essentially absent in the LA, and that the fibers were implanted more posteriorly, in the Ectorhinal cortex (Ect; FV0919).

**Figure S6.**
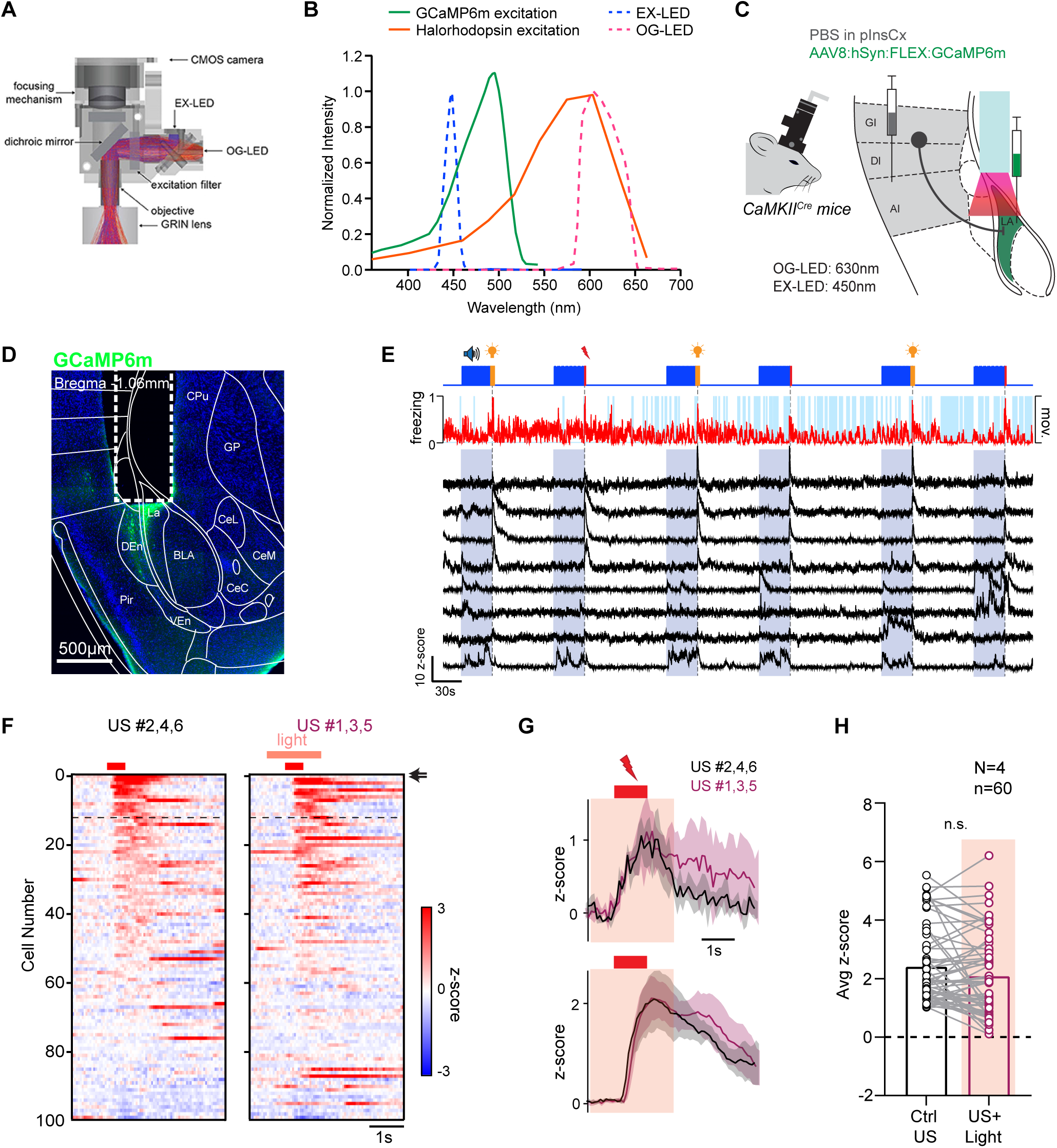
Illustration of the dual-wavelength miniature microscope approach, and control experiments for optogenetic axon silencing in *in-vivo* Ca^2+^ imaging experiments. Related to Figure 6. (A) Principle of the dual-wavelength miniature microscope used here (“nVoke”, Inscopix; image taken, with permission, from (Stamatakis et al., 2018). Two separate LEDs provide excitation for the Ca^2+^ indicator (“EX-LED”), and for activation of a longer-wavelength opsin (“OG-LED”). (B) Excitation wavelengths of the miniature microscope (“EX-LED”, and “OG-LED”, modified with permission from (Stamatakis et al., 2018), overlaid with the excitation spectrum of GCaMP6m and Halorhodopsin used here (green, and orange line; modified from Barnett et el., 2017; Chow et al., 2010). (C) Scheme of the experimental approach for the control experiments. Instead of injecting an AAV vector that drives the expression of Halorhodopsin (see Figure 6; eNpHR3.0), an equal volume of PBS was injected into the pInsCx. Scale bar, 500 µm. - Post-hoc histological validation of GCaMP6m expression and placement of the GRIN lens. Note the absence of green fluorescence in the pInsCx. This is expected, since no Halorhodopsin - eYFP was expressed in the pInsCx. (D) Ca^2+^ transients from n = 8 example imaged neurons during the training session. *Top*, movement index (red) and freezing states (light blue). (F, G) Footshock-evoked Ca^2+^ transients during no-light footshocks (*left*) and footshocks with orange light application (*right*; F), and example footshock-driven Ca^2+^ elevations from n = 2 imaged neurons for no-light footshocks and footshocks with light application (black, and purple traces, respectively). The arrows in F indicate the example neurons shown in (G). (H) Footshock-induced Ca^2+^ transient amplitude, averaged over the three no-light footshocks (*left*, black datapoints), and over the three footshocks with light application (*right*, purple data points). Light application did not induce a significant change of the footshock - driven Ca^2+^ amplitude (p = 0.074; W=-486, Wilcoxon test; data is from n = 60 US - responsive LA neurons imaged in N = 4 CamKII^Cre^ mice injected with PBS).

